# Designing Functional Dairy Food Products: Peptide-Fortification Approaches to Improve Safety, Quality, and Consumer Acceptance

**DOI:** 10.1101/2025.08.31.673363

**Authors:** Shayanti Minj, Daraksha Iram, Manish Singh Sansi, Abhishek Parashar, Kamal Gandhi, Shilpa Vij

## Abstract

This study focused on the development of functional dairy products fortified with bioactive peptides derived from Lactobacillus rhamnosus C25–fermented sheep milk and their evaluation during storage. Microencapsulated peptides were incorporated into flavoured milk and srikhand, and their antimicrobial, antioxidant, and sensory properties were monitored over refrigerated storage. Flavoured milk supplemented with free peptides exhibited strong antimicrobial activity, while encapsulated peptides provided a controlled and sustained release, maintaining functionality over six days. In srikhand, both plain and mango-flavoured formulations demonstrated enhanced biofunctional properties, with encapsulated peptides showing gradual peptide release, higher antioxidant activity, and better preservation of sensory quality over 15 days. Sheep milk used for fermentation showed good microbial quality, supporting safe peptide production. Fermentation yielded peptide fractions (<3, <5, and <10 kDa), which were analyzed using RP-HPLC and LC-MS/MS, identifying over 3,100 peptides. In silico screening predicted 34 peptides with antimicrobial potential, including cationic and amphipathic α-helical peptides primarily derived from β-casein and κ-casein. Functional assays demonstrated potent antimicrobial activity against Gram-positive and Gram-negative pathogens, particularly in the 5–10 kDa fractions. Antioxidant activity was highest in the 5 kDa retentate, indicating that medium-sized peptides contributed most to radical scavenging. Microencapsulation using sodium alginate improved peptide stability and controlled release, mitigating bitterness and preserving product acceptability. Overall, the study highlights the potential of L. rhamnosus C25–fermented sheep milk peptides as natural bioactive ingredients for functional dairy products with enhanced shelf life, antimicrobial efficacy, and antioxidant capacity.

**Graphical abstract:** 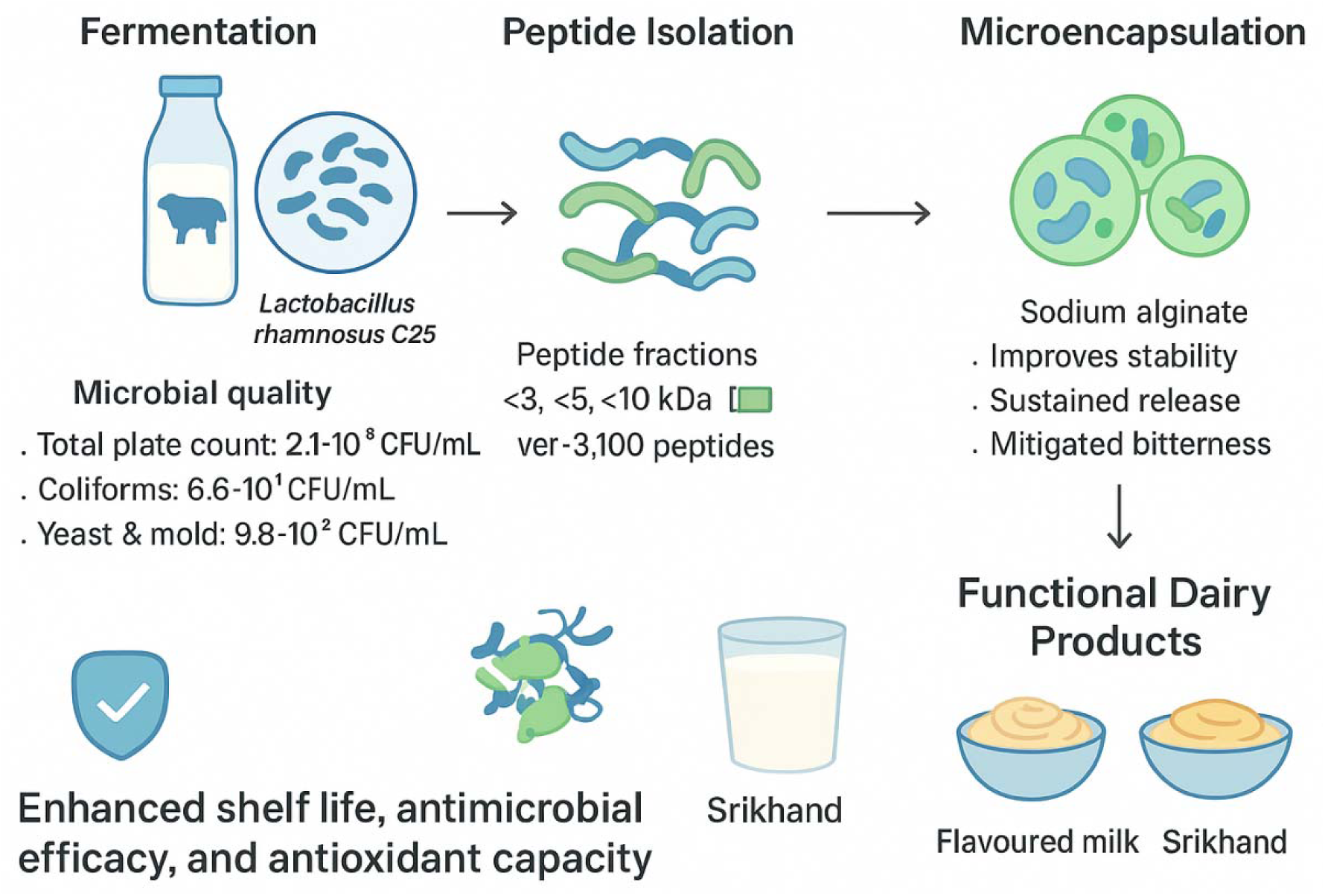

## 1.0 Introduction

Milk is considered nature’s most complete food, offering balanced nutrients and bioactive compounds that support metabolism, organ development, and disease resistance. It contains immunoglobulins, antibacterial peptides, antimicrobial proteins, oligosaccharides, and lipids that protect both infants and adults against pathogens. Milk proteins, primarily casein and whey (80:20), are key contributors to these benefits. Some proteins, such as lactoferrin, exert biological effects directly, while others release bioactive peptides through proteolysis. These peptides play crucial roles in gastrointestinal and immune system regulation, infant development, and antimicrobial defense (Schanbacher et al., 1998; Korhonen & Pihlanto, 2003; Gobbetti et al., 2007). Sheep milk is less commonly consumed as a beverage. Sheep milk is primarily used for cheese production, though in some regions it is also processed into yogurt and whey cheese (Haenlein & Wendorff, 2006). Its high nutritional value is attributed to its rich composition, with higher total solids, protein, and fat content than cow or goat milk (Haenlein, 2001; Pulina et al., 2006). Sheep milk provides high-quality protein, calcium, phosphorus, and lipids in a balanced ratio, yielding about 15% cheese compared to 10% from cow milk. It contains ∼5.8% protein, with caseins comprising 76–83% and whey proteins 17–24%. Casein fractions slightly differ from cow milk, with higher κ-CN, αs2-CN, αs1-CN, and β-CN (Moatsou et al., 2004). Sheep milk delivers about one-third more energy than cow or goat milk and is richer in beneficial fats, vitamins (including 2–3× higher Vitamin C), folate, and minerals such as calcium, iron, magnesium, phosphorus, and zinc, while being lower in sodium (Mayer & Fiechter, 2012). Two cups of milk or 93 g of sheep cheese meet daily calcium, riboflavin, and five essential amino acid requirements, while one liter provides daily protein, most essential vitamins, calcium, and phosphorus (Geerlings et al., 2006). During fermentation, casein and whey proteins release bioactive peptides (3–20 amino acids) with antimicrobial, antihypertensive, antioxidant, immunomodulatory, opioid, and mineral-binding functions.

Most milk protein bioactivities are latent and become functional only after proteolytic digestion, which releases encrypted bioactive peptides. These peptides, typically 3–20 amino acids long, regulate gastrointestinal development, immunity, microbial balance, and infant growth (Kamau et al., 2010; Korhonen, 2009). They are released during gastrointestinal digestion, fermentation by proteolytic lactic cultures, or enzymatic hydrolysis. Once absorbed, they enter circulation and exert systemic effects. Many peptides are multifunctional, displaying antimicrobial, immunomodulatory, and other biological activities (Zhang et al., 2010; Singh et al., 2014). Protein and peptide drugs, however, have short half-lives due to enzymatic degradation. Microencapsulation offers a solution by enclosing bioactives within protective coatings, enhancing stability, controlling release, and prolonging bioavailability (Singh et al., 2010; Ma, 2014). This technique is widely applied to improve shelf life, targeted delivery, and sustained release of peptides and proteins in food and pharmaceutical formulations. Milk protein-derived peptides are increasingly recognized for their health-promoting potential and application in functional foods. Extensive studies on bovine milk have identified peptides with antihypertensive, mineral-binding, antioxidant, antimicrobial, and anti-carcinogenic effects, demonstrating their physiological significance. However, research on sheep milk-derived peptides remains relatively limited, and only a few products have been commercialized. This presents a significant opportunity to explore sheep milk as a valuable source of bioactive peptides with multifunctional properties. Sheep milk contains higher protein levels and a unique casein profile, making it a promising candidate for the production of peptides with enhanced bioactivity. Such peptides could be incorporated into functional foods, nutraceuticals, and pharmaceuticals to support cardiovascular health, immune modulation, bone health, and disease prevention. Despite their potential, protein- and peptide-based therapeutics face challenges such as enzymatic degradation, short biological half-life, and poor bioavailability. Advances in delivery technologies, including microencapsulation, nanoencapsulation, and controlled-release systems, are being investigated to protect peptides during processing, enhance stability, and ensure targeted delivery in the gastrointestinal tract or systemic circulation. Future research should focus on optimizing production methods, evaluating bioavailability and efficacy through in vivo studies, and developing scalable technologies for industrial application.

## 2. Material method

### 2.1. MATERIALS

All chemical reagents utilized in this study are listed in the “Appendices” section and detailed within the manuscript. The chemicals, media, and reagents were sourced from Hi-Media Pvt. Ltd., Mumbai, and Sigma Chemical Industry, Navi Mumbai.

### 2.2. Lactic culture and Pathogenic organisms

A single, optimized lactic acid bacterial culture, *Lactobacillus rhamnosus* C25, previously isolated from cheese and characterized in our laboratory, was employed in this study. Five pathogenic bacterial strains were used as indicator organisms to assess the antimicrobial potential of the test samples. These included *Salmonella typhi* (NCTC 6017), *Enterococcus faecalis* (ATCC 27736), *Bacillus cereus* (ATCC 13061), *Listeria monocytogenes* (ATCC 15303), and *Escherichia coli* (ATCC 2592), procured from the American Type Culture Collection (ATCC) and National Collection of Type Cultures (NCTC).

### 2.3. Maintenance of lactobacillus and pathogenic cultures

The Lactobacillus rhamnosus C25 culture was grown in MRS broth at 37°C for 16–18 hours. For long-term storage, an aliquot of the overnight culture (500 µL) was mixed with an equal volume of sterilized 80% glycerol and kept at −20°C in an ultra-low temperature freezer (Blue Star, India). Before use, the culture was routinely revived by inoculating into fresh MRS broth. Additionally, a portion of the culture was propagated at 37°C in litmus milk tubes and stored in the refrigerator. All pathogenic indicator strains were maintained on Brain Heart Infusion (BHI) broth at 37°C and sub-cultured every 15 days. BHI slants were refrigerated until required, and for extended storage, 50% glycerol stocks were prepared.

### 2.4. Collection of milk samples

Sheep milk was obtained from local villages in Karnal. The milk was first screened for microbes and filtered through muslin cloth to remove debris. It was then skimmed by centrifugation at 6000 rpm for 10 minutes and subsequently sterilized by autoclaving at 121°C for 15 minutes under 15 psi. The processed milk was stored at 4°C until further use. Microbiological analysis was carried out using the pour plate method. Serial dilutions of the milk were prepared, and total bacterial count, yeast and mold count, and coliform count were determined. The protein content of the unprocessed sheep milk was measured using the Lowry protein estimation method.

### 2.5 Fermentation of sheep milk

Sheep milk was collected from villages around Karnal and initially screened for microbial contaminants. Debris was removed by filtering the milk through muslin cloth. The milk was then skimmed by centrifugation at 6000 rpm for 10 minutes and sterilized via autoclaving at 121°C for 15 minutes at 15 psi. The processed milk was stored at 4°C until further analysis. For microbiological assessment, serial dilutions of the milk were prepared, and the pour plate technique was used to determine total bacterial count, yeast and mold count, and coliform count. The protein content of the raw milk was quantified using the Lowry method.

### 2.6 Extraction of 10kda, 5kda and 3kda fractions from fermented milk

Fermented sheep milk samples were centrifuged at 6000 × g for 15 minutes at 4°C to separate bacterial cells. The resulting supernatant was first filtered through a 0.42 µm Millipore membrane and then subjected to ultrafiltration using molecular weight cut-off membranes of 3 kDa, 5 kDa, and 10 kDa (Millipore VIVASPIN) by centrifugation at 8000 × g for 15 minutes at 4°C. Fractions with molecular weights below 3 kDa, 5 kDa, and 10 kDa, as well as those above these cut-offs, were collected and stored at −20°C for further analysis.

### 2.7. Antimicrobial activity of sheep milk fermentate and its fractions (3kDa, 5kDa and 10kDa)

To evaluate the antimicrobial potential of sheep milk fermentate and its fractions against various enteric pathogens, the agar well diffusion method was employed. Freshly grown pathogens (3 hours) were mixed (70 μL) with 7 mL of molten soft agar and poured over pre-solidified nutrient agar plates. After the soft agar solidified, 6 mm wells were created using a sterile well borer and filled with 100 μL of fermentate or peptide samples. The plates were left at 4°C for 1 hour to allow sample diffusion, followed by incubation at 37°C for 12–16 hours. Zones of inhibition were measured, and a clear inhibition zone of 10 mm or greater, including the well diameter, was considered indicative of antimicrobial activity.

### 2.8. Antioxidant activity of fermented milk and its fractions (3kDa, 5kDa and 10kDa) by ABTS (2, 2 azino bis 3 ethyl benzothiazoline-6-sulphonic acid) method

This assay measures the total radical scavenging activity based on the ability of a sample to quench the stable ABTS radical within 10 minutes, with minor modifications to the method of Re et al. (1999). The ABTS working solution was prepared as described in Appendix II and subsequently diluted with phosphate-buffered saline (PBS) to achieve an absorbance of 0.7 ± 0.02 at 734 nm. A 20 µL aliquot of the sample was added to 980 µL of PBS in a cuvette and mixed for 10 seconds. The reduction in absorbance at 734 nm was recorded over 10 minutes at 10-second intervals using a spectrophotometer. Results were expressed as Trolox equivalent antioxidant capacity (TEAC). All measurements were performed in triplicate. For the standard curve, 12.5 mg of Trolox (molecular weight 250.9) was dissolved in 10 mL ethanol to prepare a 5 mM stock solution. This stock was diluted with distilled water to generate concentrations ranging from 0 to 1000 µM. The standard curve was constructed by plotting Trolox concentration (µM) on the X-axis versus % inhibition on the Y-axis. Experiments were performed in triplicate.

### 2.10. Microencapsulation of bioactive peptides for the preparation of flavoured milk and Srikhand

The 30 kDa peptide fraction was chosen for microencapsulation as it includes smaller bioactive peptides of approximately 10 kDa, 5 kDa, and 3 kDa, which are known to possess notable antimicrobial activity. Encapsulating the entire 30 kDa fraction helps protect these bioactive peptides, enhancing their stability and allowing controlled release during processing and storage. The microencapsulated 30 kDa fraction was subsequently incorporated into flavored milk, producing a functional dairy product with potential health benefits.

### 2.11. Microencapsulation of 30 kDa

Fermented sheep milk samples were centrifuged at 8000 × g for 15 minutes at 4°C to separate bacterial cells. The resulting supernatant was first filtered through a 0.42 µm Millipore membrane and then through an ultrafiltration membrane with a 30 kDa molecular weight cut-off. The fraction below 30 kDa was collected and used for microencapsulation. Sodium alginate (2% w/v) served as the wall material for preparing the microcapsules.

**Figure 1.**
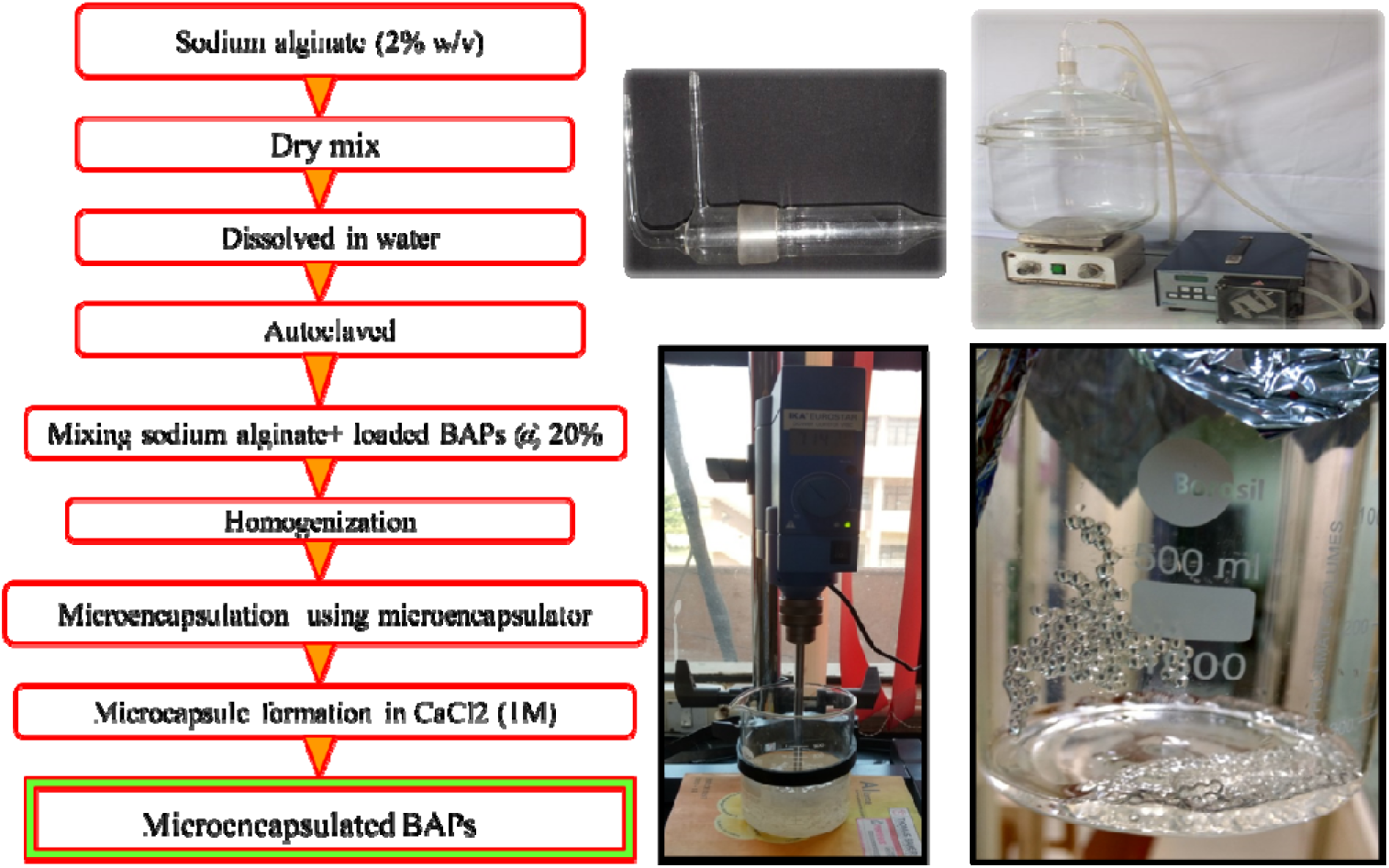
Microencapsulation through two fluid nozzle air atomization method

### 2.12. Preparation of core-coat mixture

The core–coat mixture was prepared by dissolving the polymer in distilled water using a magnetic stirrer (Labco Instruments, New Delhi, India) and sterilizing it by autoclaving. Specifically, 4 g of sodium alginate was added to 200 mL of distilled water, and the peptide sample was incorporated at 20% (v/v). A 0.1 M calcium chloride solution was prepared by dissolving 220 g of calcium chloride in 10 L of distilled water and autoclaving; this solution was used as the hardening agent.

### 2.13. Preparation of microcapsules

A concentric two-fluid glass nozzle with air atomization was employed for microencapsulation. The molten core–coat mixture was delivered via a peristaltic pump (Ravel Hiteks Pvt. Ltd., Chennai, India) at a controlled flow rate into the air-atomizing nozzle and sprayed under pressurized air into a reaction vessel containing 3 liters of 0.1 M calcium chloride solution, with continuous magnetic stirring at 1000 rpm for 30 minutes. The distance between the nozzle tip and the liquid surface was maintained at approximately 12 inches. The inner nozzle had a 1 mm orifice, with a 1.3 mm annular gap between the inner and outer nozzle. Pressurized air disrupted the matrix fluid into fine droplets, which fell into the calcium chloride solution. The divalent calcium ions cross-linked with sodium alginate, forming calcium–alginate microcapsules through ionotropic gelation. The resulting microcapsules were collected by sieving through a Buchner funnel, washed twice with sterile distilled water, and stored for further studies.

### 2.14. Encapsulation Efficiency

Encapsulation efficiency was determined by comparing the total peptide content in the final encapsulated product with the peptide concentration in the initial feed solution.

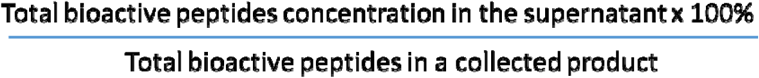

### 2.15. Pathogen Inhibition Assay

The growth inhibition assay was performed following a modified protocol of Balasubramaniam et al. (2011). The antimicrobial activity of peptides encapsulated in the alginate system was assessed by counting colony-forming units (CFUs). Bacillus cereus ATCC 13061 was cultured in BHI broth at 30°C for 24 hours. Three sets of flasks, each containing 50 mL of BHI broth, were inoculated with 100 µL of the bacterial culture. One flask set served as the control (C), while unencapsulated peptides were added to the second set (2%, 8000 AU) and encapsulated peptides were added to the third set (15% capsules, 8000 AU). Samples from each flask were taken at 0, 2, 4, 8, and 12 hours, serially diluted, plated on BHI agar, and CFUs were enumerated.

### 2.16. Preparation of flavoured milk and srikhand

Sheep milk-derived bioactive peptides were utilized as functional food ingredients. After microencapsulation, these peptide fractions were incorporated into flavored milk to develop a functional dairy product. The milk was first preheated and supplemented with microcapsules at 7%, then bottled and stored for further analysis. The standardized preparation process involved adjusting milk to 2% fat and preheating to 35–40°C, followed by the addition of sugar (10%), color and flavouring agents (0.03%), and microcapsules (7%). The mixture was then pasteurized, cooled to 5°C, and packaged for storage at 5°C. This controlled process ensures product safety, stability, and maintenance of flavor during storage (Kumar et al., 2013).

### 2.17. Physicochemical analysis of flavoured milk

Freshly prepared flavored milk samples were evaluated for physicochemical properties, including pH, fat, sugar, and total protein content.

#### 2.17.1. pH

The pH of the flavored milk samples was measured using a pH meter (SDFCL, India), which was initially calibrated with standard buffers of pH 4.0 and 7.0 at 20.0 ± 0.1°C.

#### 2.17.2. Total fat

The fat content of flavored milk samples was determined using the Mojonnier Extraction Method (AOAC, 2000). Ten milliliters of milk sample were placed into an extraction tube, followed by 1.25 mL of ammonia (sp. gr. 0.8974) and mixed thoroughly. Next, 10 mL of ethyl alcohol was added and mixed, followed by 25 mL of peroxide-free diethyl ether. The tube was stoppered and shaken vigorously for about one minute. Then, 25 mL of petroleum ether (boiling range 40–60°C) was added and mixed for approximately 30 seconds. The mixture was allowed to stand until the upper ether layer separated and became clear, after which it was decanted into a suitable container. The extraction tube was washed with a small amount of ether, and the washings were added to the collected ether layer. The remaining liquid in the tube was washed twice with 15 mL of each solvent, and the washings were combined with the ether layer. The ether was then completely evaporated. The flask containing the extracted fat was dried in an air oven at 102 ± 2°C for 2 hours, cooled in a desiccator, and weighed. It was heated again for 30 minutes, cooled, and reweighed. Fat was carefully washed out from the flask using petroleum ether, leaving the insoluble residue behind. The flask was dried and weighed again. The difference in weights represented the amount of fat extracted from the flavored milk. A blank determination using the reagents was performed to calculate the corrected fat content.

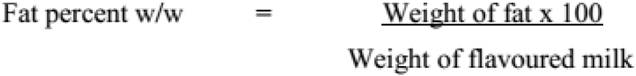

#### 2.17.3. Sugar

A 40 g portion of a well-mixed sample was placed in a 100 mL beaker, and 50 mL of hot water was added. The mixture was transferred to a 250 mL volumetric flask, and the volume was adjusted to 150 mL. After cooling to room temperature, 5 mL of dilute ammonia solution was added and allowed to stand for 15 minutes. An equivalent amount of dilute acetic acid was then added to neutralize the ammonia. The mixture was further combined with 12.5 mL of zinc acetate solution and 12.5 mL of potassium ferrocyanide solution, and the final volume was adjusted to the 250 mL mark. The solution was allowed to settle and then filtered; the filtrate was designated as solution B1.

From solution B1, 50 mL was pipetted into a 100 mL volumetric flask, followed by the addition of 5 mL of concentrated hydrochloric acid. The mixture was heated in a water bath at 65–68°C for 5 minutes, cooled, and neutralized with sodium hydroxide solution. This resulting solution was labeled A1. Solutions B1 and A1 were diluted to obtain volumes (A2 and B2) such that 15–50 mL would react with 10 mL of Fehling’s solution. The diluted solutions were transferred to 50 mL burettes. In a 250 mL conical flask, 10 mL of mixed Fehling’s solution was heated for 1–2 minutes, and the prepared solutions were added gradually from the burettes until the blue color of the indicator disappeared. One milliliter of methylene blue indicator was then added, and the titration was completed by carefully adding the solutions in small increments.

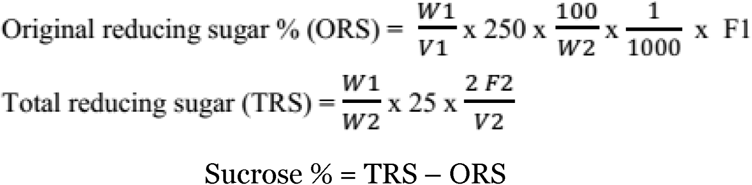

Where,

W1 = weight in milligram of sucrose corresponding to 10ml of Fehling’s solution

W2 = weight in gram of sample taken

F2 = dilution factor for solution A2

V2 = volume in ml of solution A2 corresponding to 10ml of Fehling’s solution

F1 = dilution factor for solution B2

V1 = volume in ml of solution B2 corresponding to 10ml of Fehling’s solution

#### 2.17.4. Total protein

The protein content of the flavored milk samples was measured using the Lowry method (Lowry et al., 1951).

### 2.18. Storage study of flavoured milk

The flavored milk samples were divided into three groups: a control (FC), a sample supplemented with unencapsulated peptides (FF), and a sample containing microcapsules (FE). All samples were stored at 4°C for up to one week. Analyses of biofunctional properties were conducted at 2-day intervals, specifically on days 0, 2, 4, and 6, to evaluate antimicrobial, antioxidant, and antihypertensive activities.

#### 2.18.1. Sensory evaluation of flavoured milk

The product’s sensory attributes were evaluated by a panel of eight untrained participants.

### 2.19. Preparation of double emulsion

Water-in-oil-in-water (W/O/W) double emulsions were prepared by first dispersing the peptide samples in the water phase and combining them with an oil phase to form the primary emulsion. This primary emulsion was then mixed with an aqueous phase containing a water-soluble stabilizer (pectin) to form the secondary emulsion. The double emulsions were produced using high-speed homogenization (Sharma, 2015).

**Figure 2.**
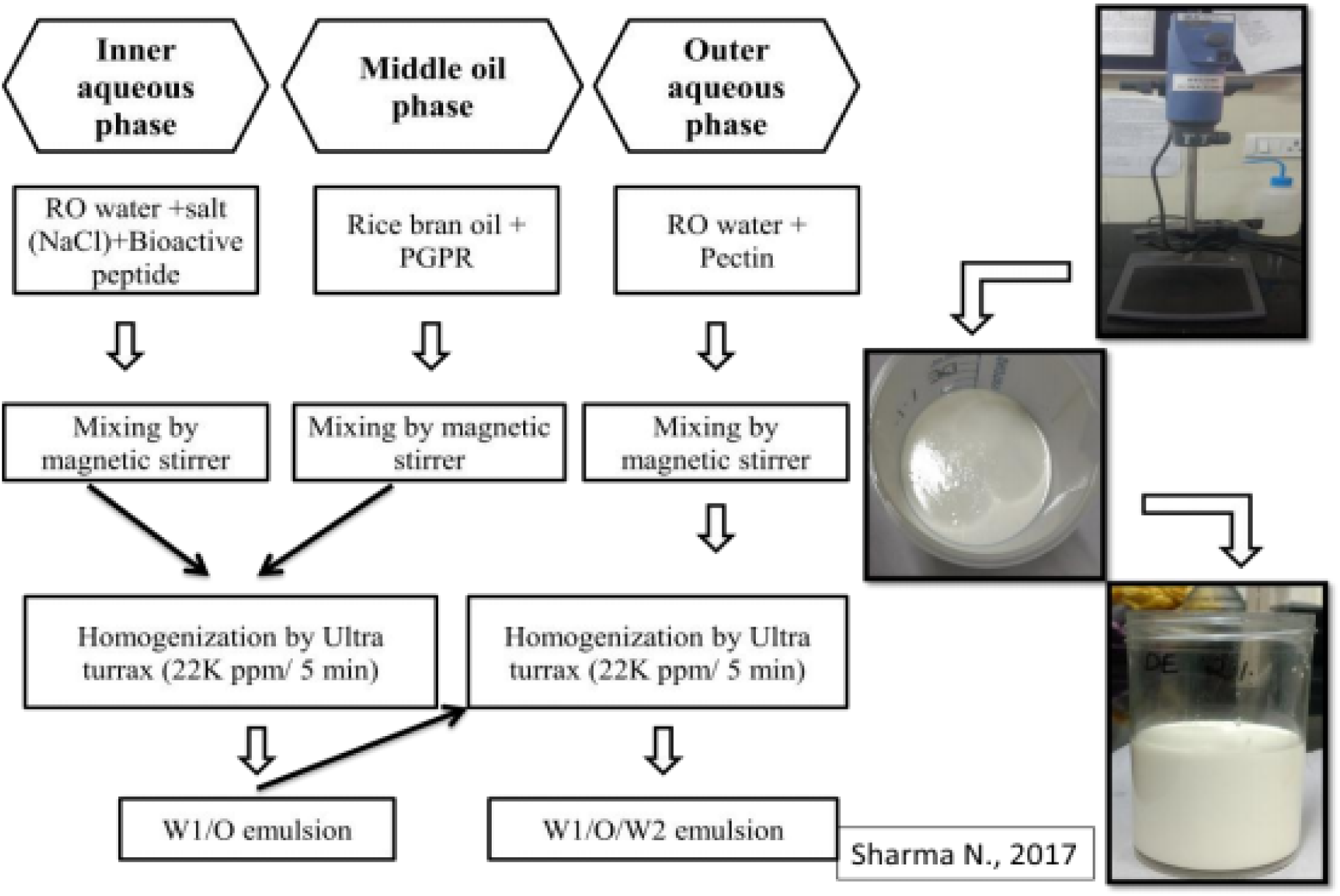
Preparation of double emulsion

#### 2.19.1. Addition of double emulsion in flavoured milk

The double emulsion was incorporated into flavored milk at 7%. No further analysis was performed due to phase separation of the emulsion.

**Figure 3.**
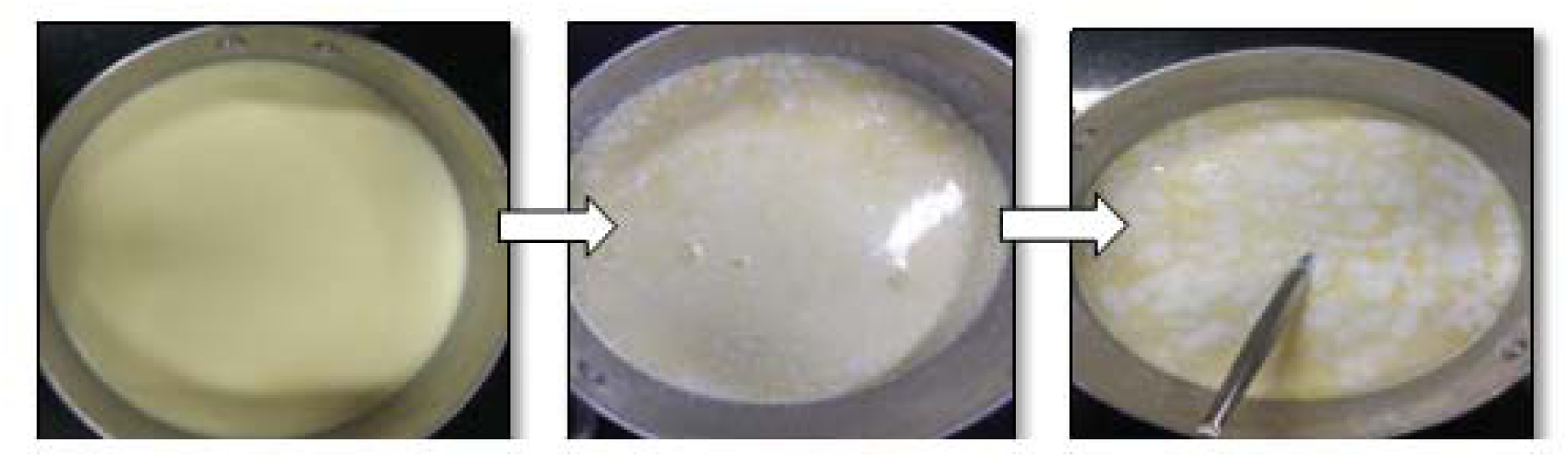
Addition of double emulsion in flavoured milk

### 2.20. Preparation of Srikhand with Double Emulsion

Milk was first adjusted to the desired fat and SNF content. The standardized milk was preheated at 90°C for 10 minutes and then cooled to 37°C. It was inoculated with starter culture at 2% (v/v) and incubated at 30°C for 16 hours to form curd. The whey was drained overnight to produce chakka (concentrated curd). The chakka was mixed with 30% sugar, 2% double emulsion, and 0.02% cardamom. For the mango variant, 15% mango pulp was added. The mixture was homogenized to achieve uniform texture, then packaged into 20 g cups, sealed, and stored under refrigeration until further analysis.

**Figure 4.**
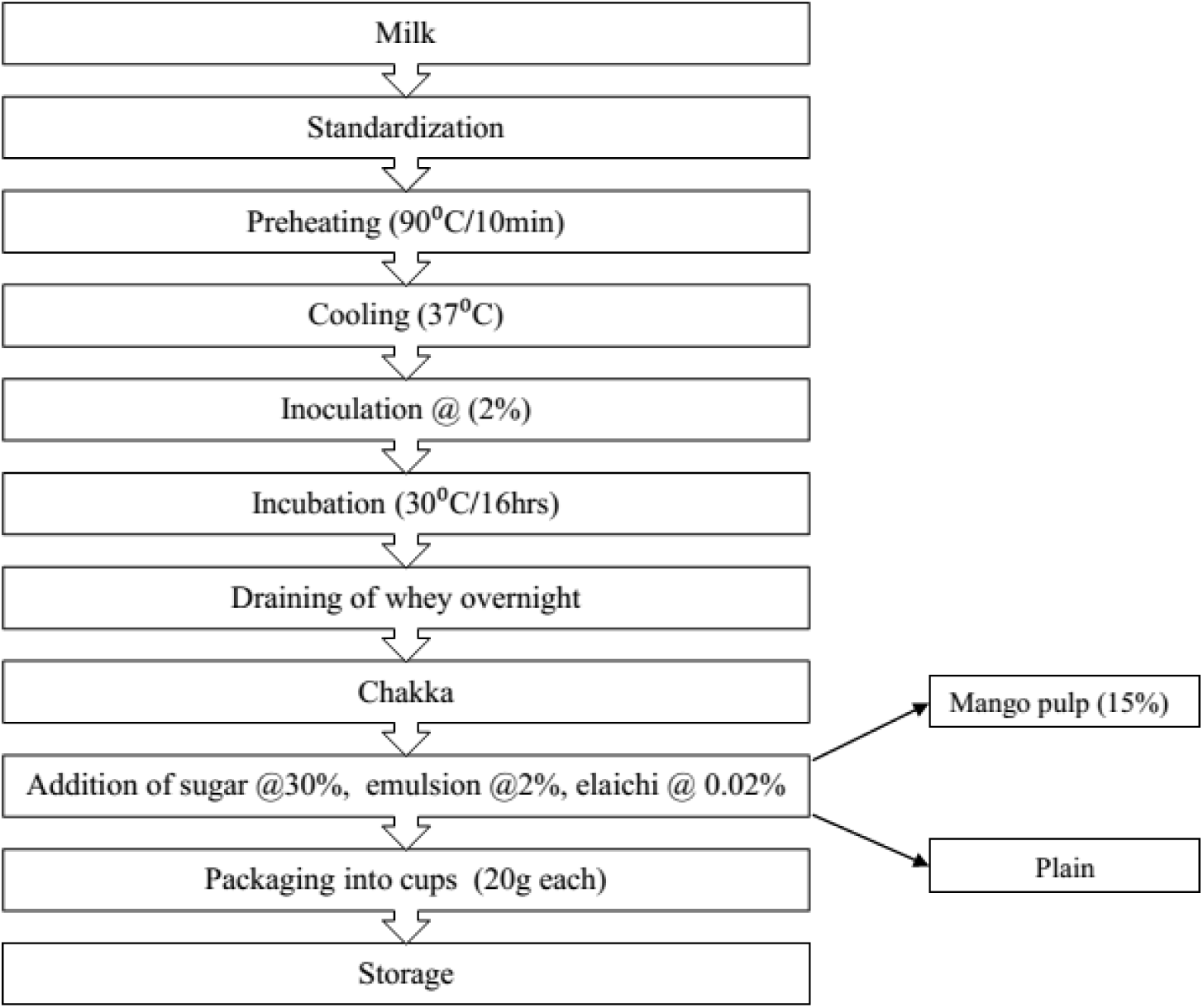
Addition of double emulsion in srikhand

#### 2.20.1. Physicochemical analysis of srikhand

Freshly prepared plain srikhand samples were evaluated for physicochemical properties, including pH, fat, sugar, and total protein.

### 2.21. Storage study of srikhand

Mango srikhand samples were divided into three groups: a control (MC), a sample supplemented with unencapsulated peptides (MF), and a sample containing encapsulated peptides (ME). Similarly, plain srikhand was divided into control (PC), unencapsulated peptide (PF), and encapsulated peptide (PE) groups. All samples were stored at 4°C for up to 14 days. Analyses of biofunctional properties were conducted at 3-day intervals, specifically on days 0, 3, 6, 9, 12, and 15, to assess antimicrobial, antioxidant activities.

#### 2.21.1. Microbial analysis and Sensory evaluation of srikhand

Srikhand samples were monitored for total microbial count on Nutrient Agar at 3-day intervals throughout the 15-day storage period (days 0, 3, 6, 9, 12, and 15). The product’s sensory characteristics were evaluated by a panel of seven judges, and overall acceptability was recorded.

### 2.22. RP-HPLC analysis of sheep milk bioactive peptides

Sheep milk proteins and peptides obtained after fermentation were analyzed by RP-HPLC using a C18 column, following the method of Vincenzetti et al. (2008). The mobile phases were: (A) 0.1% (v/v) trifluoroacetic acid (TFA) in deionized water and (B) 0.1% (v/v) TFA in 100% acetonitrile. A linear gradient was applied as follows: 100% solvent A for 5 min; solvent B gradient 0–5 min, 0%; 5–10 min, 0–50%; 10–30 min, 50–65%; 30–35 min, 65–100%; 35–40 min, 100%; 40–45 min, 100–0% solvent B. The flow rate was maintained at 0.6 mL/min, and the eluent was monitored at 214 nm using a UV detector (Najafian et al., 2014).

### 2.23. Identification and sequencing of milk derived bioactive peptides by mass spectrometry (LC-MS/MS)

Peptides from different fractions (<10 kDa, <5 kDa, <3 kDa) were sequenced using nano-flow LC-MS/MS (Captive Spray–Maxis HD Impact II QTOF, Bruker, Germany). Separation was achieved on a C18 nano-trap column (Bruker Magic C18AQ, 0.1 × 150 mm, 3 µm particle size, 200 Å pore size). Mobile phase A consisted of 0.1% (v/v) formic acid in Optima LC-MS grade water, while mobile phase B contained 0.1% (v/v) formic acid and 2% (v/v) water in Optima LC-MS grade acetonitrile. A gradient elution was performed at 5 µL/min: 2% B for 2 min, increasing to 20% B at 46 min, 45% B at 61 min, then 95% B over 2 min, held for 5 min, and finally returned to 2% B for 12 min, for a total run time of 84 min. Analytes were detected in positive ion mode using LC-MS over an m/z range of 130–2200 at a 2 Hz sampling rate. LC-MS/MS detection was performed in positive ion mode at 5 Hz, covering m/z 50–2000, allowing for up to 10 precursors per cycle. All reagents used were of LC-MS grade.

### 2.24. *In silico* prediction of potential antimicrobial peptides (AMPs)

Peptides identified from fermented sheep milk via LC-MS/MS were screened using the PeptideRanker web server (http://bioware.ucd.ie/). Peptides with a bioactivity score greater than 0.70 were considered potential bioactive candidates (Mooney et al., 2012). Further in silico analysis of antimicrobial peptides (AMPs) was performed using two web-based prediction tools, iAMPpred and the ADAM server (Meher et al., 2016; Yu, 2015).

### 2.25. Prediction of physicochemical properties of antimicrobial peptides

Peptide properties, including net charge, theoretical isoelectric point, molecular weight, and GRAVY (grand average of hydropathicity) index, were calculated using the ProtParam web tool (https://web.expasy.org/cgi-bin/protparam/ProtParam). Peptide bioactivity was predicted using PeptideRanker, and toxicity was assessed with the ToxinPred tool (http://crdd.osdd.net/raghava/toxinpred/). The distribution of hydrophobic and hydrophilic residues in the peptides was analyzed using HeliQuest (Gautier et al., 2008). Additionally, the three-dimensional structures of the peptides were modeled using the UCSF Chimera software (Gasteiger et al., 2005; Gupta et al., 2013; Gautier et al., 2008; Pettersen et al., 2004).

### 2.26. Peptide profiling of fermented sheep milk

Bioactive antimicrobial peptide profiles were generated for αs1-casein, κ-casein, and β-casein proteins of sheep milk using the UniProtKB database. Peptides exhibiting antimicrobial properties were then compared with those identified through the LC-MS/MS analysis.

## 3. Result and discussion: Purification of cultures

### 3.3.1. Microscopic examination

The purity and morphology of *Lactobacillus rhamnosus* C25 and pathogenic strains were evaluated using Gram staining. Microscopic examination under oil immersion (100×) revealed that *L. rhamnosus* C25 appeared as small Gram-positive rods. In contrast, the pathogenic isolates showed variation in Gram reaction (positive or negative) as well as in cell shape and arrangement (rods or cocci). Catalase testing confirmed that all pathogenic cultures were catalase-positive, whereas *L. rhamnosus* C25 was catalase-negative.

### 3.2 Microbiological analysis of milk

Sheep milk was procured from local villages of Karnal and subjected to microbiological quality assessment, including total plate count (TPC), coliform count, and yeast and mold enumeration. The results showed a TPC of 2.1×10□ CFU/mL, coliform count of 6.6×10¹ CFU/mL, and yeast and mold count of 9.8×10² CFU/mL, indicating good microbial quality of the milk. Previous studies have reported higher microbial loads in raw milk. For instance, Benkerroum et al. (2003) observed mean counts of aerobic bacteria (6.2×10□ CFU/mL), enterococci (2.9×10□ CFU/mL), coliforms (1.6×10□ CFU/mL), LAB (7.0×10□ CFU/mL), yeasts (1.0×10□ CFU/mL), *Staphylococcus aureus* (3.8×10□ CFU/mL), and sulphite-reducing clostridia (6 CFU/mL). Similarly, Houser et al. (2008) reported considerably higher standard plate counts (9.7×10□ CFU/mL) and coliform counts (3.2×10□ CFU/mL) in raw milk compared to bulk tank milk.

### 3.3 Production of bioactive peptides from sheep milk

Bioactive peptide production was carried out by fermenting camel and goat milk with Lactobacillus rhamnosus C25 in 1000 mL conical flasks under optimized conditions (Arun, 2016), with and without shaking. A rapid decline in pH was observed within the first 6 h of fermentation, which stabilized after 24 h. Following fermentation, the supernatant was collected, sequentially filtered through 0.45 μm and 0.22 μm filters, and stored at 4°C. The filtrate was then fractionated using molecular weight cut-off (MWCO) membranes to obtain peptide fractions of 10, 5, and 3 kDa.

### 3.4. Characterization of sheep milk fermented bioactive peptides using RP-HPLC

Sheep milk proteins and fermented peptide fractions (<3 kDa, <5 kDa, and <10 kDa) were analyzed using RP-HPLC, and their chromatograms are presented in Figure 1. In sheep milk, distinct peaks corresponding to α-, β-, and κ-caseins were observed at retention times of approximately 5, 6.2, and 9.7 min, respectively, along with a small α-lactalbumin peak at 6.3 min (Fig. 1). The <10 kDa peptide fraction displayed several peaks between 5 and 25 min, with most appearing between 10 and 20 min. Similarly, the <5 kDa fraction exhibited peaks within 5–22 min, while the <3 kDa fraction showed well-defined peaks in the 5–22.5 min range, with a major concentration of peaks between 5 and 15 min. These peaks were subjected to sequencing and identification using LC-MS/MS to determine their bioactive peptide sequences.

**Figure 1.**
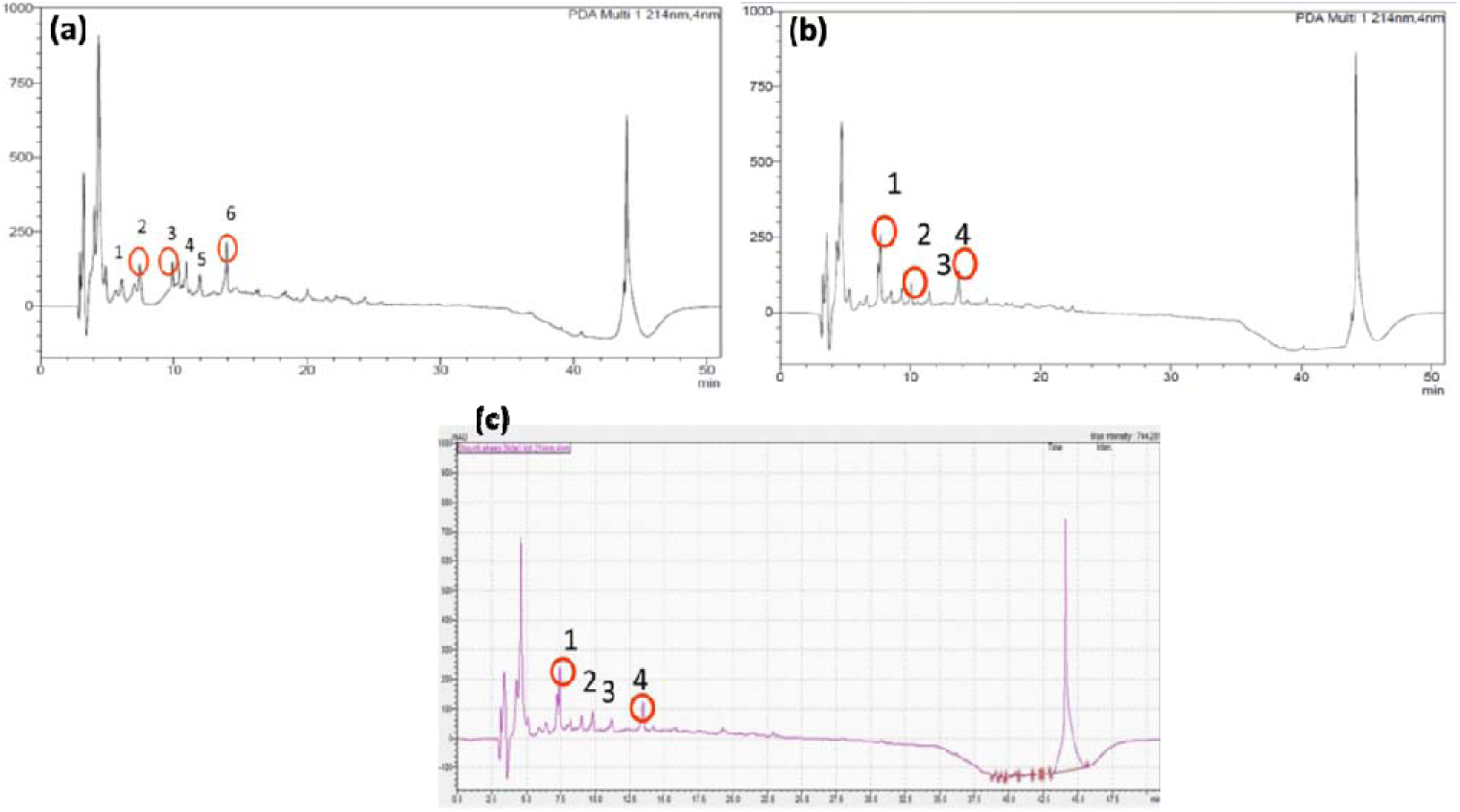
RP-HPLC Chromatogram of (a) 10kDa (b) 5kDa and (c) 3kDa fraction of sheep milk

### 3.5. Antimicrobial activity of fermented sheep milk peptides

The antimicrobial activity of sheep milk fermentate and its peptide fractions was evaluated using the agar well diffusion method against five indicator pathogens. The highest inhibition was recorded against *B. cereus* (10 ± 0.57 mm), followed by *E. coli, E. faecalis, S. typhi*, and *L. monocytogenes*. Compared to the crude fermentate, all peptide fractions (10, 5, 3 kDa) exhibited enhanced antimicrobial effects. The 10 kDa permeate showed the strongest activity, with inhibition zones of 28.33 ± 0.66 mm (*B. cereus*), 22.66 ± 0.88 mm (*E. coli*), and similar trends for other pathogens. The 5 kDa retentate displayed the next highest activity, whereas the 3 kDa fractions showed relatively lower inhibition. These results suggest that the most potent bioactive peptides are smaller than 10 kDa but larger than 5 kDa showed in figure 2.

The antimicrobial potential of milk-derived peptides is largely attributed to their interaction with microbial cell membranes. These peptides are typically small (fewer than 50 amino acids), cationic, and amphiphilic, which facilitates their electrostatic interaction with negatively charged bacterial surfaces. This interaction is considered the initial step leading to membrane destabilization, pore formation, and ultimately, bacterial cell lysis (Benkerroum, 2010). In some cases, the peptides are also capable of penetrating the membrane to target intracellular components, further enhancing their antimicrobial effects (Matsuzaki, 1999). Bacterial membranes are rich in negatively charged phospholipids, which increases their affinity for cationic peptides. The amphiphilic nature of these peptides, possessing both hydrophobic and hydrophilic regions, allows them to integrate into lipid bilayers while remaining soluble in aqueous environments (Power et al., 2013; Kim & Wijesekara, 2010). Experimental studies have demonstrated this mechanism in action. For instance, ovine milk caseinate hydrolyzed with protease from Bacillus sp. P7 displayed antibacterial activity against Bacillus cereus (9.3 mm) and *Corynebacterium fimi* (11.5 mm), whereas other tested pathogens remained resistant (Correa et al., 2011). Similarly, ovine αs2-casein digested with pepsin for 30 min exhibited inhibitory activity against both Gram-positive and Gram-negative bacteria (Lopez-Exposito et al., 2006). These findings highlight the potential of milk-derived peptides as promising natural alternatives to conventional antibiotics.

**Figure 2.**
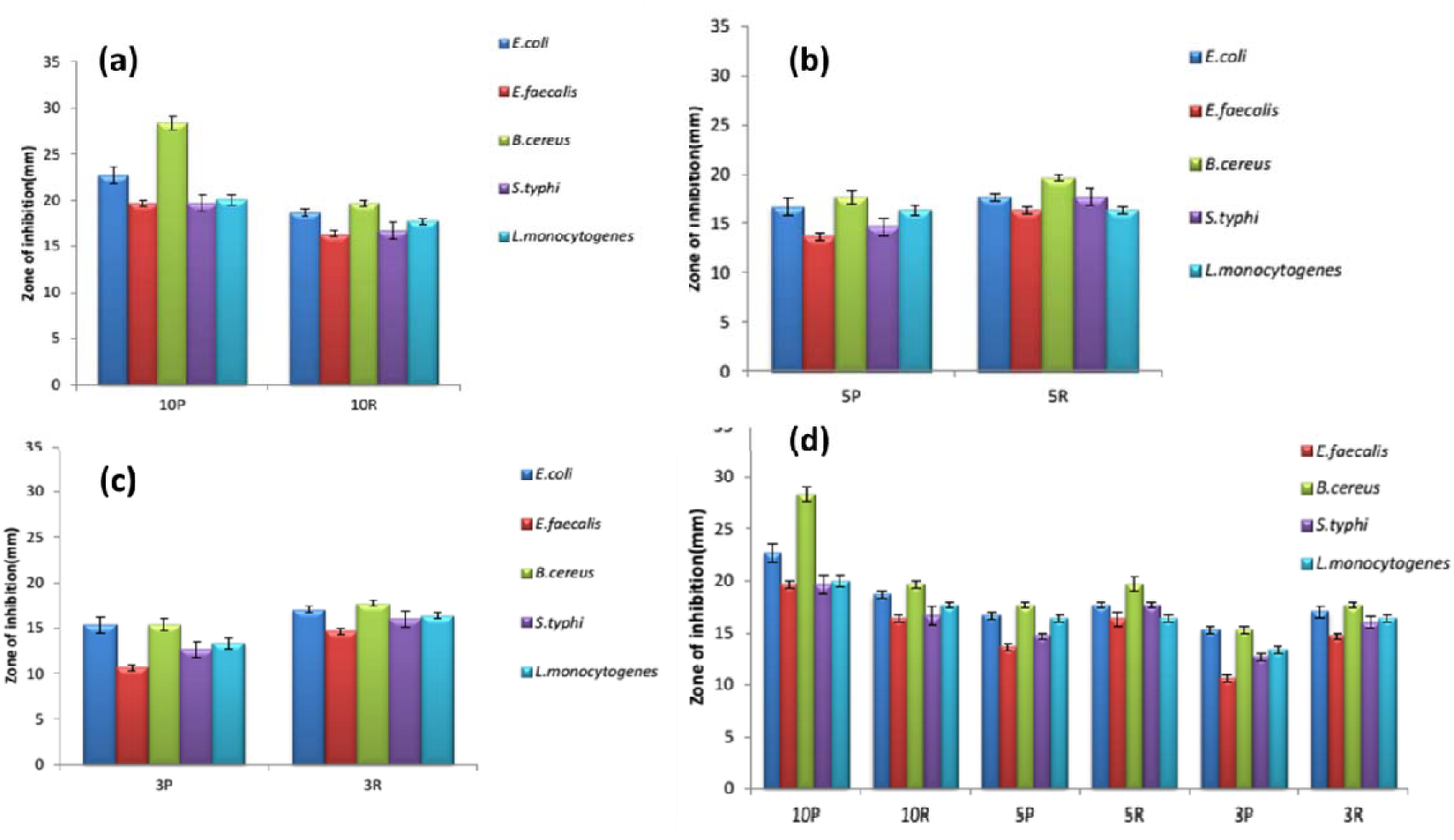
(a) Antimicrobial activity of sheep milk peptides <10 to >10 kDa MWCO (b) Antimicrobial activity of sheep milk <5 to 5-10 kDa MWCO (c) Antimicrobial activity of sheep milk <3 to 3-5 kDa MWCO (d) Comparative antimicrobial activity of sheep milk peptides of <3 to >10 kDa MWCO

### 3.6. Antioxidant activity of sheep milk peptides

The antioxidant capacity using the ABTS assay was measured with Trolox as the standard, a vitamin E analogue with strong radical scavenging ability. Results were expressed as Trolox Equivalent Antioxidant Capacity (TEAC). In this assay, the ABTS•+ radical cation is reduced to a colorless form in the presence of antioxidants capable of donating hydrogen or breaking radical chain reactions. This method is suitable for evaluating both hydrophilic and lipophilic antioxidants. In the present study, antioxidant activity was observed in all peptide fractions derived from sheep milk fermented with Lactobacillus rhamnosus C25, including fractions ranging from <3 kDa to >10 kDa. The TEAC values were as follows: 10 kDa permeate – 2230 µmol/L, 10 kDa retentate – 1909.8 µmol/L, 5 kDa permeate – 2195.5 µmol/L, 5 kDa retentate – 2279.98 µmol/L, 3 kDa permeate – 1853.29 µmol/L, and 3 kDa retentate – 1990.09 µmol/L. Among these, the highest antioxidant capacity was recorded in the 5 kDa retentate fraction, followed by the 10 kDa permeate fraction showed in figure 3. These findings suggest that the most potent antioxidant peptides are smaller than 10 kDa but larger than 5 kDa. Previous studies support these results. Correa et al. (2011) reported that hydrolysis of ovine caseinate with Bacillus sp. P7 protease enhanced the antioxidant capacity of the resulting hydrolysates up to 2 h of hydrolysis, with no further increase beyond that point. This indicates that enzymatic treatment liberates previously inactive peptides embedded within the casein sequence, contributing to antioxidant potential. Gomez-Ruiz et al. (2008) also demonstrated that the antioxidant activity of ovine casein is influenced by its primary amino acid sequence and structural conformation. Hydrolysis of intact ovine casein and its fractions with pepsin, trypsin, and chymotrypsin produced peptides with significant ABTS radical-scavenging activity (Gomez-Ruiz et al., 2008). Similarly, water-soluble peptide extracts from ovine and goat cheese-like systems exhibited antioxidant potential when evaluated using the ABTS assay (Silva et al., 2006).

**Figure 3.**
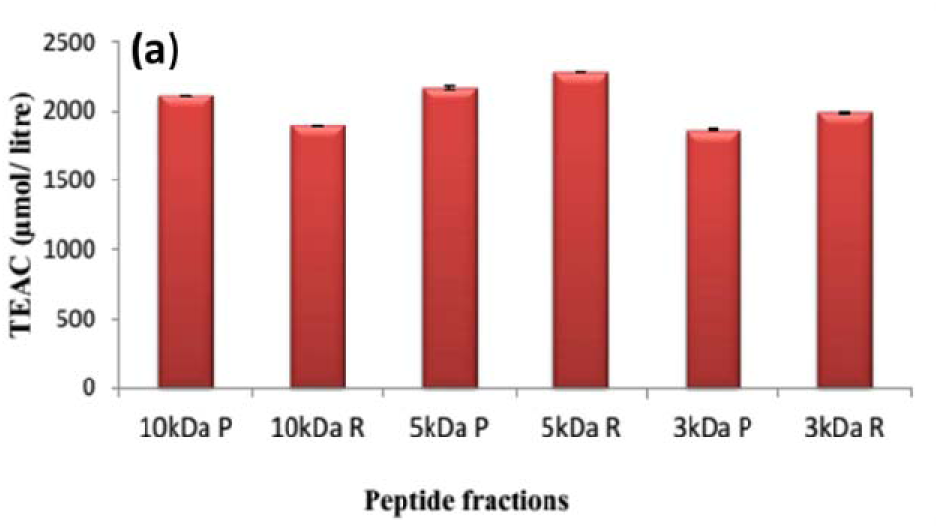
Antioxidant activities of sheep milk protein fractions (10kDa, 5kDa, and 3kDa) permeate and retentate

### 3.7. Microencapsulation of sheep milk bioactive peptides

Consumers today increasingly seek foods that go beyond basic nutrition, expecting them to offer added health benefits. Food protein hydrolysates and bioactive peptides have gained attention as potential functional food ingredients due to their physiological benefits. However, their commercial use remains limited, as they often face challenges such as low bioavailability, bitterness, hygroscopic nature, and potential interactions with the food matrix. To address these limitations, microencapsulation is widely recognized as an effective strategy to enhance peptide stability, improve bioavailability, and mask undesirable sensory properties. In this study, fermented sheep milk samples were subjected to ultrafiltration using a membrane with a molecular weight cut-off of 30,000 Da. The permeate containing peptides smaller than 30 kDa was collected and utilized for microencapsulation. Sodium alginate (2% w/v) served as the wall material to prepare the microcapsules.

### 3.8. Antimicrobial activity of 30kDa peptide fraction

The antimicrobial potential of the 30 kDa permeate and retentate peptide fractions of sheep milk was evaluated using the agar well diffusion assay against five indicator pathogens. Both fractions demonstrated inhibitory activity against Gram-positive and Gram-negative bacteria. The zone of inhibition ranged between 17.3 ± 0.33 mm and 22.33 ± 0.33 mm (including well diameter). Among the pathogens tested, the 30 kDa permeate exhibited the highest inhibition against *Bacillus cereus* (22.33 ± 0.33 mm), followed by *Escherichia coli* (21 ± 0.5 mm), *Enterococcus faecalis* (20 ± 0.5 mm), *Salmonella typhi* (20.66 ± 0.33 mm), and *Listeria monocytogenes* (19.66 ± 0.66 mm). Similarly, the 30 kDa retentate showed the maximum inhibition against *B. cereus* (19.66 ± 0.33 mm), followed by *E. coli* (18.33 ± 0.33 mm), *E. faecalis* (16.66 ± 0.33 mm), *S. typhi* (18 ± 0.57 mm), and *L. monocytogenes* (17.3 ± 0.33 mm) showed in figure 4. Since the permeate fraction displayed stronger antimicrobial activity against all test organisms compared to the retentate, the 30 kDa permeate peptides were selected for subsequent microencapsulation studies.

**Figure 4.**
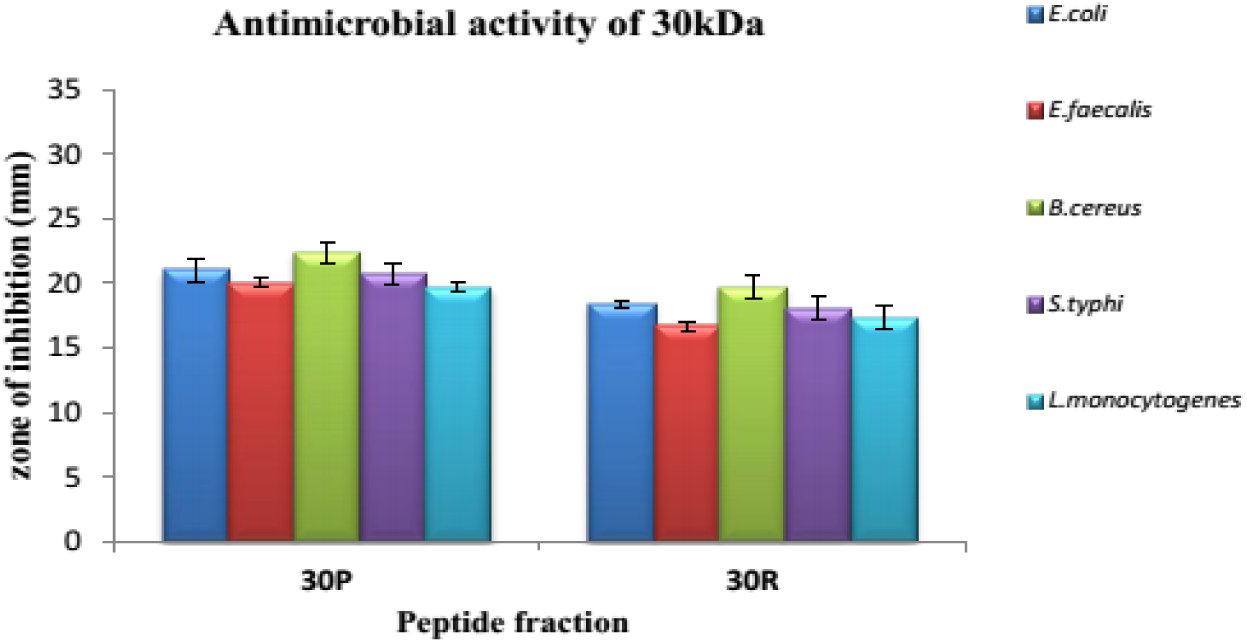
Antimicrobial activity of sheep milk 30kDa peptide fraction

### 3.9. Encapsulation efficiency

The 30 kDa permeate fraction of sheep milk was encapsulated at a 20:80 (v/v) ratio using 2% (w/v) sodium alginate as the encapsulating material, following the procedure described by Goyal (2013). Similar findings were reported by Goyal (2013), where bacteriocin encapsulation using sodium alginate resulted in an encapsulation efficiency of 35.63%. In the present study, encapsulation efficiency was determined using the Lowry protein estimation method and was found to be 37.88%, suggesting a moderate degree of peptide entrapment within the alginate matrix.

### 3.10. Pathogen inhibition assay

Growth inhibition was assessed by modifying the method described by Balasubramaniam et al. (2011). The antimicrobial potential of the 30 kDa permeate fraction encapsulated in sodium alginate was tested against Bacillus cereus by counting the colony-forming units (CFU). B. cereus cultures were grown in BHI broth at 30 °C for 24 h, and 100 µl of inoculum was added to three sets of flasks, each containing 50 ml of fresh BHI broth. One flask served as the control (C), the second received 2% of the unencapsulated peptide fraction, and the third was supplemented with 7% of the encapsulated peptides. Samples were collected at 0, 2, 4, 8, 12, and 16 h, serially diluted, and plated on BHI agar to determine viable counts.

Figure 4a, illustrates the log CFU/ml data. The control group showed a continuous increase in bacterial count, reaching nearly four log cycles. In contrast, both unencapsulated and encapsulated peptide treatments exhibited a linear reduction in bacterial counts by up to two log cycles within the first 8 h. After 8 h, a slight rise of approximately one log cycle was observed, which could be attributed to the bacteriostatic effect of the peptides, resulting in slower bacterial growth. Another possible reason could be a reduced affinity of the peptides for the bacterial cell membrane over time, leading to diminished antimicrobial activity (Pellegrini, 2003).

**Figure 5.**
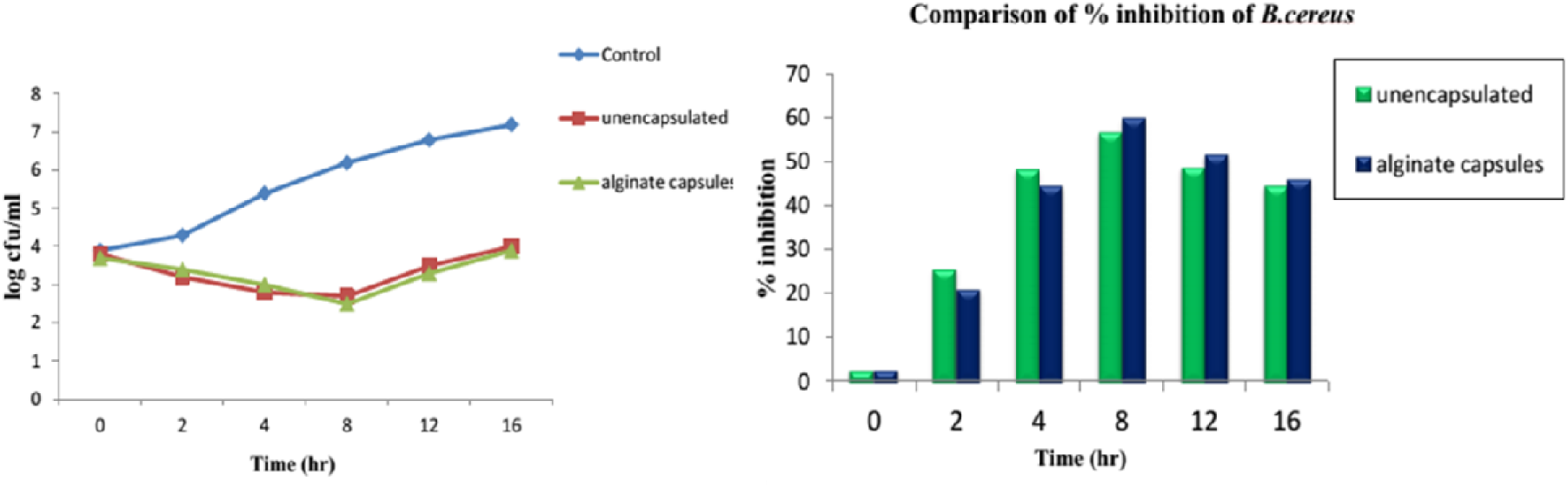
(a) Effect of unencapsulated free peptides and alginate microcapsules on growth of *B. cereus* (b) Percentage inhibition of *B. cereus* by unencapsulated free peptides and alginate microcapsulated peptides

To evaluate the comparative effect of free and microencapsulated peptide fractions, the percentage of growth inhibition was calculated. Figure 4b, illustrates the inhibitory effect of both treatments against *B. cereus*. At the initial time point (0 h), no significant difference in inhibition was observed between unencapsulated and encapsulated peptides. After 4 h, alginate microcapsules achieved a 44.44% reduction in colony count, whereas free peptides showed slightly higher inhibition at 48.14%. Interestingly, after 8 h, microencapsulated peptides demonstrated superior antibacterial activity, achieving 59.67% inhibition compared to 56.45% for free peptides. By 16 h, both treatments showed a slight decline in inhibition, with microcapsules reducing bacterial counts by 45.83% and free peptides by 44.44%.

### 3.11. Preparation of flavoured milk supplemented with bioactive peptide ingredients

In recent years, food proteins have emerged as a primary source of bioactive peptides due to their beneficial effects on various physiological processes. Incorporating such peptides into food systems through microencapsulation offers an effective approach to enhance their stability, functionality, and health-promoting properties. The food industry has increasingly adopted encapsulation techniques to address challenges associated with the rising demand for functional ingredients (Champagne & Fustier, 2007). Several studies have explored the production of bioactive peptides from milk, evaluating their functional roles and potential as ingredients in functional foods that positively influence consumer health (Albenzio et al., 2017). In the present work, microencapsulated sheep milk-derived bioactive peptides were incorporated into flavoured milk to evaluate their functional potential. The flavoured milk was divided into three groups: control (FC), milk supplemented with unencapsulated free peptides (FF), and milk containing microencapsulated peptides (FE). All samples were stored under refrigeration (4 °C) for one week, and biofunctional properties were analyzed at two-day intervals.

### 3.12. Physico-chemical parameters of flavoured milk

The physicochemical analysis of flavoured milk included the assessment of fat, protein, pH, and sugar content. Result showed that the control sample contained 2.06% fat, 3.62% protein, and 9.7% sugar, with a pH value of 6.47.

### 3.13. Analysis of biofunctional properties of flavoured milk

#### 3.13.1 Antimicrobial activity of flavoured milk

The antimicrobial properties of the three flavoured milk formulations were evaluated during storage at 4 °C for up to six days. The inhibition zones recorded throughout the storage period ranged from 6.3 ± 0.33 mm to 15.33 ± 0.33 mm, including the 6 mm well diameter. On day 0, the control sample exhibited no noticeable antimicrobial effect against the tested pathogens, except for *B. cereus* (6.33 ± 0.33 mm) and *E. coli* (6 mm), which could be attributed to the presence of inactive milk proteins. The flavoured milk supplemented with free peptides (FF) demonstrated the strongest antimicrobial activity against both Gram-positive and Gram-negative bacteria, showing the highest inhibition against *B. cereus* (13.33 ± 0.33 mm), followed by *E. coli* (12.6 ± 0.33 mm). Samples containing encapsulated peptides (FE) also displayed antimicrobial effects, though to a lesser extent than the unencapsulated peptides, likely due to the slower release of bioactive compounds from the alginate matrix. The highest inhibition in FE samples was observed against *B. cereus* (7 ± 0.5 mm), followed by *E. coli* (6.6 ± 0.33 mm) showed in figure 6. After six days of storage, FF samples continued to show a higher level of activity, with maximum inhibition recorded against *B. cereus* (15.33 ± 0.33 mm) and *E. coli* (15.0 mm), possibly due to structural rearrangements of the bioactive peptides enhancing their antimicrobial potential (Jenssen et al., 2006). No significant variation in antimicrobial activity was observed in FE samples over the storage period, indicating that peptide release remained slow and steady. The consistently lower activity in FE samples compared with FF suggests that incomplete release of the encapsulated peptides possibly caused by sedimentation may have limited their bioavailability.

**Figure 6.**
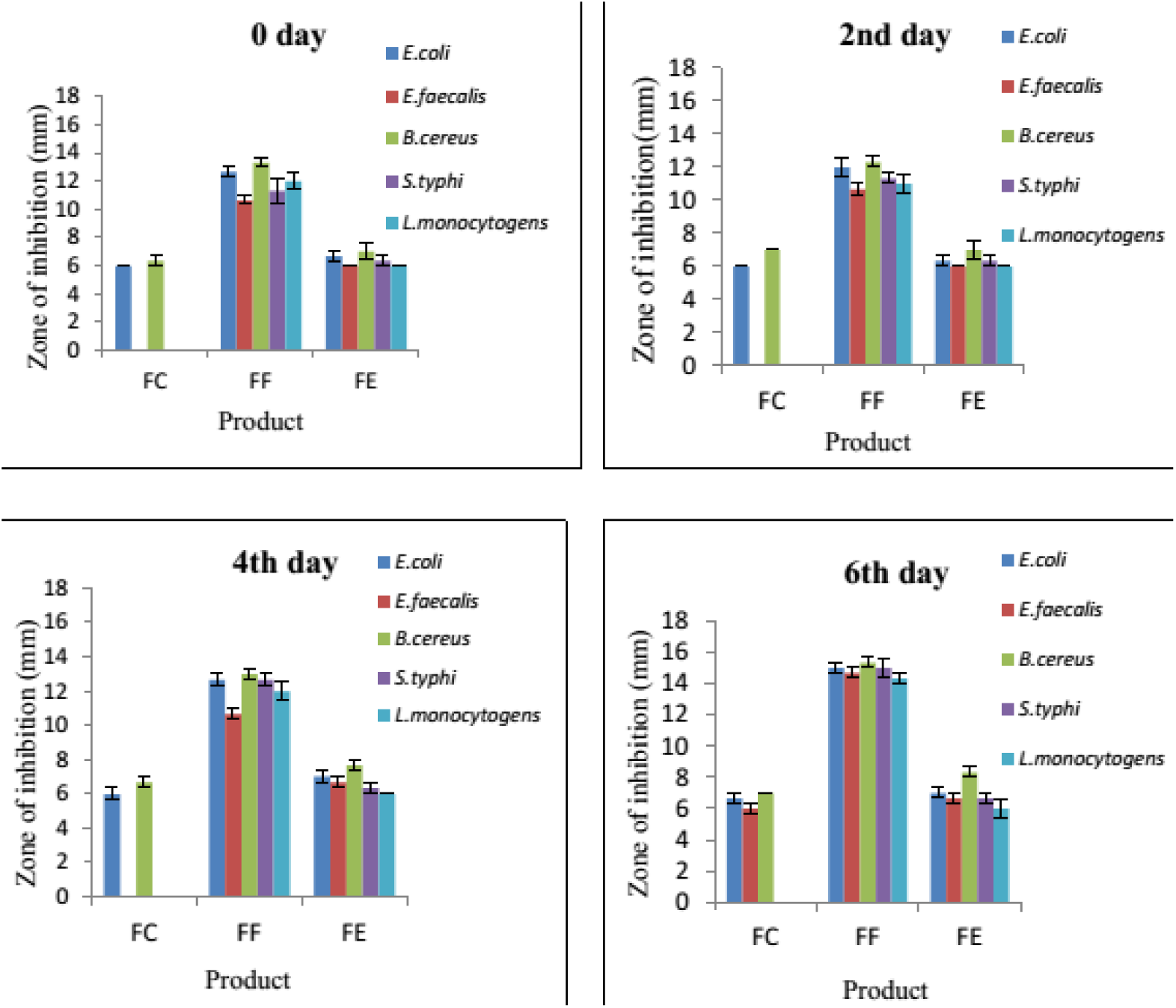
Changes in the antimicrobial activity of flavoured milk during storage *F-Flavoured milk, C-Control, F-Unencapsulated Free peptides, E-Microencapsulated peptides

### 3.14. Antioxidant activity of flavoured milk

The antioxidant potential of the three flavoured milk formulations was assessed during storage at 4 °C for up to six days. On day 0, the sample containing unencapsulated free peptides exhibited the highest antioxidant activity (976.33 ± 0.88 µmol/L TEAC), which was significantly (P < 0.05) greater than both the control (443 ± 0.2 µmol/L TEAC) and the sample with microencapsulated peptides (464.33 ± 1.2 µmol/L TEAC). The lower activity in the encapsulated sample may be attributed to the slow and gradual release of peptides from the microcapsules. No significant change in antioxidant activity was observed in the free peptide and control samples during the first two days of storage, showing 980.33 ± 0.88 and 443.36 ± 0.33 µmol/L TEAC, respectively. In contrast, a slight increase was noted in the encapsulated peptide sample (501 ± 0.88 µmol/L TEAC), suggesting progressive peptide release over time. By the end of the six-day storage period, a moderate increase in antioxidant activity was observed in the unencapsulated peptide sample (1081.67 ± 4.09 µmol/L TEAC), whereas a slight decline was recorded in the control (440.65 ± 4.09 µmol/L TEAC) showed in figure 7. The encapsulated peptide sample maintained a relatively stable activity (517.33 ± 0.33 µmol/L TEAC), indicating controlled and sustained release of bioactive peptides throughout storage.

**Figure 7.**
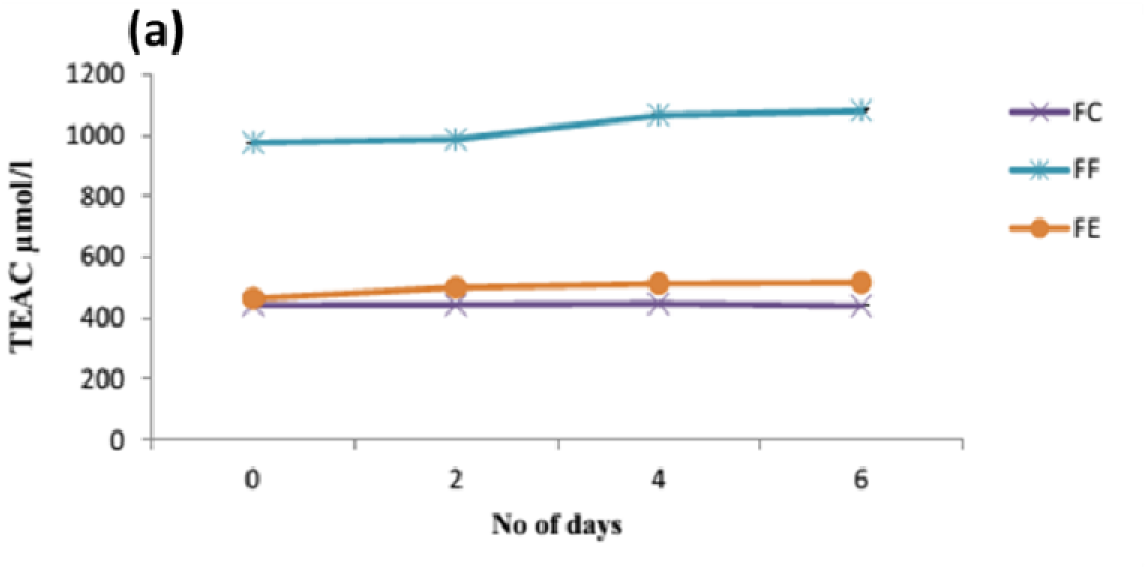
Changes in the antioxidant activity of flavoured milk during storage, *F-Flavoured milk, C-Control, F-Unencapsulated Free peptides, E-Microencapsulated peptides

### 3.15. Sensory evaluation of flavoured milk

Sensory evaluation plays a crucial role in determining the consumer acceptance of any newly developed product. It allows developers to assess whether the product meets consumer preferences based on key attributes such as flavor, body and texture, color and appearance, and acidity. In this study, sensory analysis of the flavoured milk samples was conducted for one week using a panel of eight untrained members. On day 1, the control sample received the highest overall acceptability score (87.25 ± 0.36 out of 100), followed by the sample containing unencapsulated free peptides (83.75 ± 0.72). In contrast, the sample containing microencapsulated peptides scored significantly lower (58.37 ± 0.7) showed in figure 8, likely due to sedimentation of the microcapsules and a resulting coarse mouth feel. Sensory evaluation was repeated on the fifth day of storage. The control sample again received the highest score (71.87 ± 0.65), with the free peptide sample showing no significant difference (70.5 ± 0.85), possibly due to effective masking of peptide bitterness by the added mango flavor. However, the microencapsulated peptide sample showed a further decline in acceptability (57 ± 0.7), likely caused by swelling of alginate microcapsules during storage, which could have increased sedimentation and negatively affected texture. Based on these results, sodium alginate microencapsulated peptides were deemed unsuitable for flavoured milk development, as their presence negatively impacted sensory quality.

**Figure 8.**
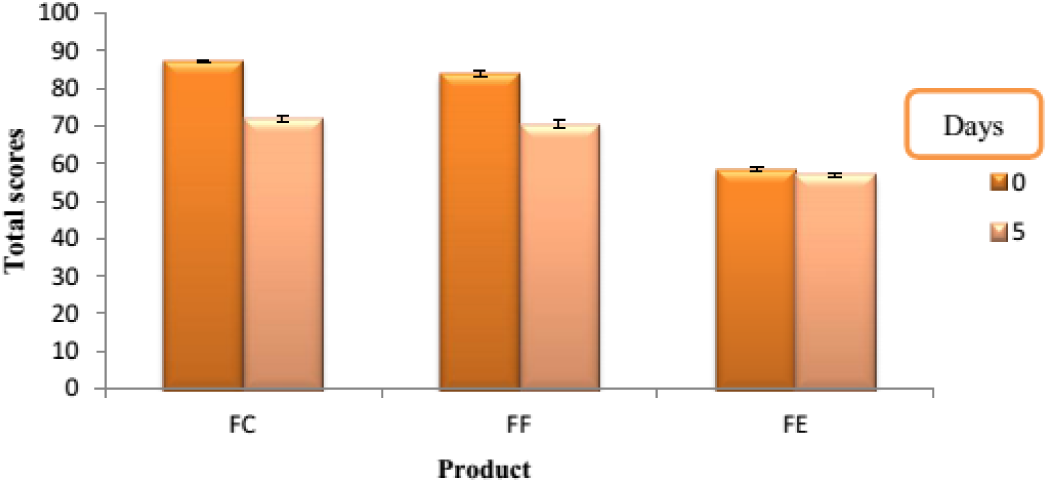
Sensory evaluation of flavoured milk during storage *F-Flavoured milk, C-Control, F-Unencapsulated Free peptides, E-Microencapsulated peptides

### 3.16 Preparation of double emulsion

Significant progress has been made in recent years in the development of structured delivery systems designed for the encapsulation of nutraceuticals and bioactive food ingredients. Multiple types of delivery systems are now available, each offering unique benefits and limitations depending on the intended application. Among these, lipid-based systems prepared using emulsion technology are the most widely utilized in the food sector. Oil-in-water emulsions, in particular, are extensively applied across several industries, including pharmaceuticals, cosmetics, healthcare products, agrochemicals, and functional foods, due to their ability to act as efficient carriers for bioactive components. In the present work, a double emulsion (water-in-oil-in-water) system was formulated. The process involved dissolving 30 kDa sheep milk peptides (aqueous phase) into an oil phase to create the primary emulsion. This was then further emulsified with a pectin-containing aqueous phase to form the secondary emulsion showed in figure 9. The double emulsion was prepared using a high-speed homogenization technique and subsequently incorporated into flavoured milk to evaluate the functional potential of the peptides. However, phase separation was observed in the final product, which might be attributed to density differences and phase incompatibility. Double emulsion technology offers several advantages for functional food development, including reduced active compound concentration, protection of sensitive bioactives, and controlled release properties. It also holds potential for applications such as salt and fat reduction while maintaining product quality and sensory appeal (Garti, 1997; Sapei et al., 2012). Despite these benefits, commercial use of double emulsions in food products remains limited due to challenges with long-term stability. Ongoing research aims to enhance the stability of double emulsions and facilitate their successful integration into functional food systems (Benichou et al., 2004; Cofrades et al., 2013).

### 3.17. Preparation of srikhand

In this research, sheep milk bioactive peptides encapsulated using a double emulsion system were incorporated into srikhand as functional ingredients. The double emulsion blended well with the srikhand matrix, allowing further evaluation of the biofunctional properties of the peptides in a real food system. Two types of srikhand were prepared plain and mango-flavoured (with 15% mango pulp). Each type was divided into three groups: a control sample, a sample with unencapsulated free peptides, and a sample containing encapsulated peptides. All samples were stored under refrigeration (4 °C) for up to two weeks, and analyses of biofunctional activities were conducted at three-day intervals throughout the storage period.

### 3.18. Physico-chemical parameters of srikhand

The physicochemical analysis of srikhand included evaluation of fat, protein, pH, and sugar content. The results indicated a pH of 5.14, fat content of 5.63%, protein content of 10.2%, and sugar content of 70.8%.

### 3.19. Biofunctional properties of srikhand

#### 3.19.1. Antimicrobial activity of srikhand

The antimicrobial potential of both plain and mango srikhand was evaluated during storage at refrigeration temperature (4 °C) for up to two weeks. The inhibition zones throughout the storage period ranged from 6 ± 0.33 mm to 15.6 ± 0.33 mm for plain srikhand, and from 6 mm to 15.6 ± 0.33 mm for mango srikhand (including the 6 mm well diameter). All samples exhibited antimicrobial activity (Figure 10). On day 0, control samples of both plain and mango srikhand showed the lowest inhibition, with the highest activity observed against *Bacillus cereus* (6.66 ± 0.33 mm), followed by inhibition against *Listeria monocytogenes* (6 ± 0.33 mm). No significant increase in antimicrobial activity was observed in the control samples during storage (days 0, 3, 6, 9, 12, and 15). In contrast, srikhand samples fortified with unencapsulated free peptides and encapsulated peptides demonstrated considerably higher antimicrobial activity on day 0, with maximum inhibition observed against *B. cereus* (12.66 ± 0.33 mm and 12.62 ± 0.33 mm, respectively).

**Figure 10.**
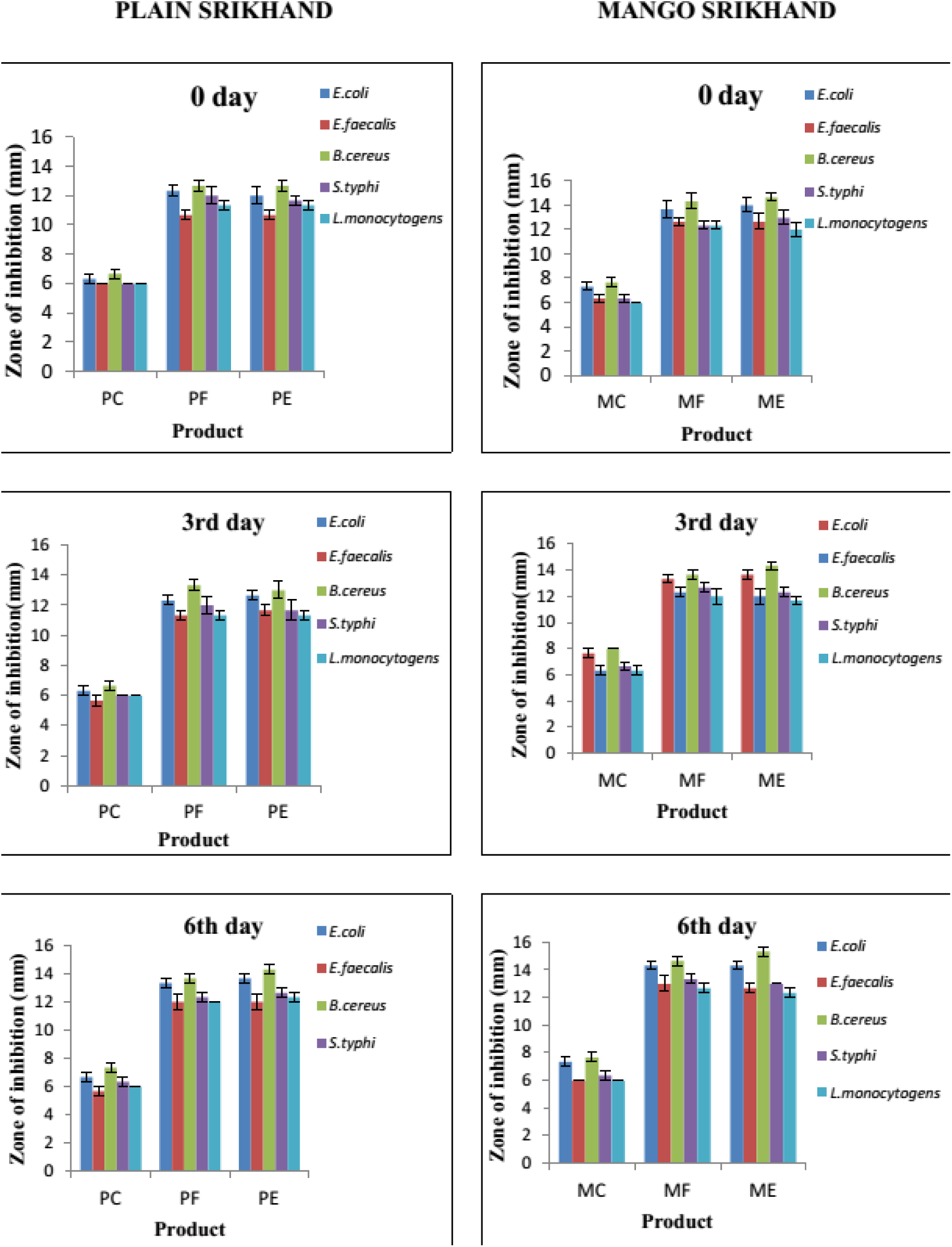
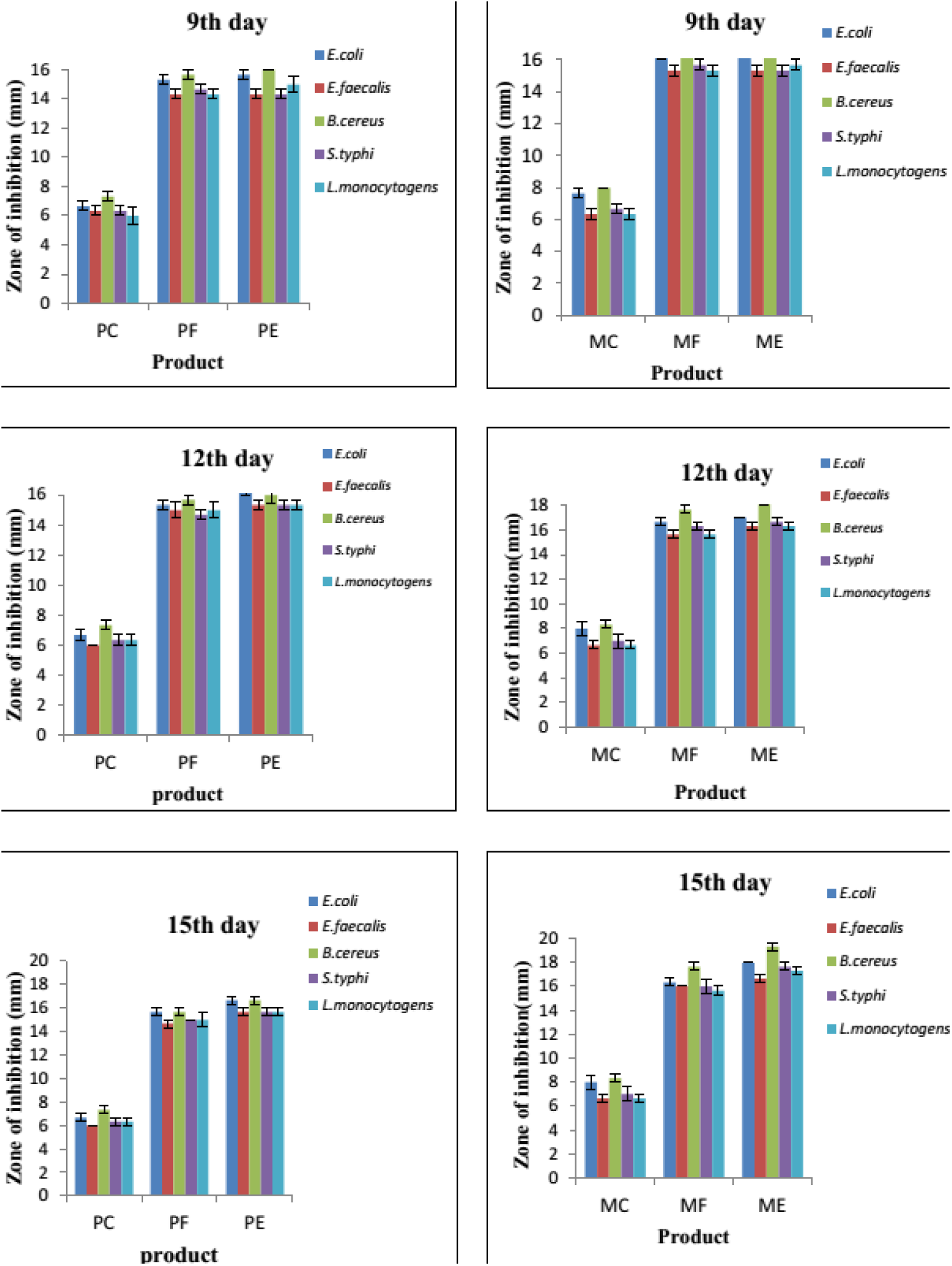
Changes in antimicrobial activity of plain and mango srikhand during storage *M-Mango, P-Plain, C-Control, F-Unencapsulated Free peptides, E-Encapsulated peptides

The samples also exhibited inhibitory activity against *L. monocytogenes*, with inhibition zones measuring 11.33 ± 0.33 mm and 11.3 ± 0.33 mm for unencapsulated and encapsulated peptides, respectively. Over the storage period (days 0, 3, 6, 9, 12, and 15), no significant difference in antimicrobial activity was observed between the samples containing unencapsulated and encapsulated peptides at the initial stage. However, by the end of storage, the samples with encapsulated peptides demonstrated a slight increase in inhibition against all tested pathogens, likely due to the sustained release of peptides from the double emulsion system. In contrast, the antimicrobial activity of samples containing unencapsulated peptides remained unchanged, possibly due to a reduced binding affinity of cationic peptides toward the pathogens, allowing some bacterial growth. Furthermore, no significant difference in antimicrobial activity was observed between plain and mango srikhand across all treatments (control, unencapsulated peptides, and encapsulated peptides) throughout the storage period.

### 3.20. Antioxidant activity of srikhand

The antioxidant potential of plain and mango srikhand was evaluated over 15 days of refrigerated storage at 4 °C (Figure 11). Significant effects (P < 0.05) of treatment (control, unencapsulated peptides, and encapsulated peptides), storage duration, srikhand type, and their interactions were observed on antioxidant activity. On day 0, control samples exhibited the lowest antioxidant activity, recording 720.3 ± 0.35 µmol/L TEAC in plain srikhand (PC) and 900.3 ± 0.58 µmol/L TEAC in mango srikhand (MC). In contrast, samples supplemented with unencapsulated peptides showed higher activity (976.33 ± 0.35 µmol/L TEAC in PF and 1289.11 ± 0.60 µmol/L TEAC in MF), while those with encapsulated peptides exhibited similar or slightly higher values (976 ± 0.57 µmol/L TEAC in PE and 1291.28 ± 0.38 µmol/L TEAC in ME). During storage (days 0, 3, 6, 9, 12, and 15), both peptide-enriched treatments demonstrated a significant (P < 0.05) increase in antioxidant capacity. By day 15, mango srikhand with unencapsulated and encapsulated peptides reached 1701.66 ± 1.2 µmol/L TEAC and 1900.44 ± 0.23 µmol/L TEAC, respectively, while plain srikhand reached 1080.66 ± 1.2 µmol/L TEAC (PF) and 1135.2 ± 1.3 µmol/L TEAC (PE). In contrast, control samples showed no notable change, maintaining values around 804.43 ± 1.2 µmol/L TEAC (PC) and 994.06 ± 0.52 µmol/L TEAC (MC). Interestingly, encapsulated peptide treatments displayed a slightly greater improvement in activity toward the end of storage, likely attributable to the controlled and gradual release of peptides, whereas free peptide activity remained relatively stable, possibly due to the reduced bioactivity of peptides over time. When comparing the two varieties, mango srikhand consistently exhibited significantly higher antioxidant activity than plain srikhand across all treatments throughout storage, which may be attributed to the presence of bioactive compounds from mango pulp and rice bran oil in the emulsion, known for their antioxidative properties (Ribeiro et al., 2007).

**Figure 11.**
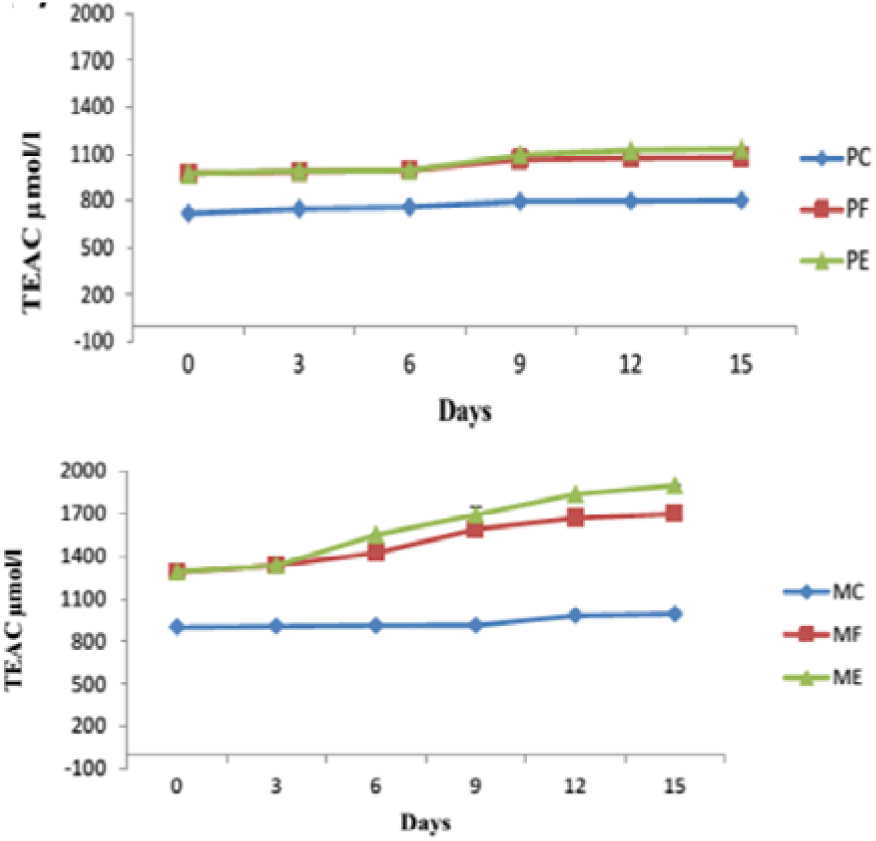
Changes in the antioxidant activity of plain and mango srikhand with added peptide ingredients during storage; *M-Mango, P-Plain, C-Control, F-Unencapsulated Free peptides, E-Encapsulated peptides.

### 3.20. Microbial analysis of srikhand

The microbiological assessment of both plain and mango srikhand samples was performed to evaluate the total microbial load and possible contamination during storage. Total viable counts were determined using nutrient agar up to 15 days of refrigerated storage. The pour plate method was employed after preparing serial dilutions, followed by incubation at 37 °C. Significant effects (P<0.05) were noted for treatment (control, free peptides, and encapsulated peptides), storage period (0, 3, 6, 9, 12, and 15 days) showed in figure 12, srikhand type, and their interactions. On day 0, microbial counts were nearly identical among all treatments in plain srikhand (8.70, 8.73, and 8.71 log CFU/ml) and mango srikhand (8.61, 8.63, and 8.60 log CFU/ml). Over the storage period, control samples showed a slight increase in microbial counts, reaching 8.96 ± 0.02 log CFU/ml in plain and 8.77 ± 0.08 log CFU/ml in mango srikhand. In contrast, samples containing free peptides exhibited a small but significant reduction in microbial load by day 15 (8.59 ± 0.01 log CFU/ml in plain and 8.42 ± 0.02 log CFU/ml in mango), likely due to the antimicrobial action of the peptides against lactic acid bacteria. Encapsulated peptide-treated samples also showed a slight decrease in microbial counts (8.50 log CFU/ml in plain and 8.32 log CFU/ml in mango). However, no significant difference was observed between the free peptide and encapsulated peptide treatments. Overall, the interaction between srikhand type, treatment, and storage duration was not statistically significant (P>0.05), and microbial counts remained comparable between plain and mango srikhand throughout storage.

**Figure 12.**
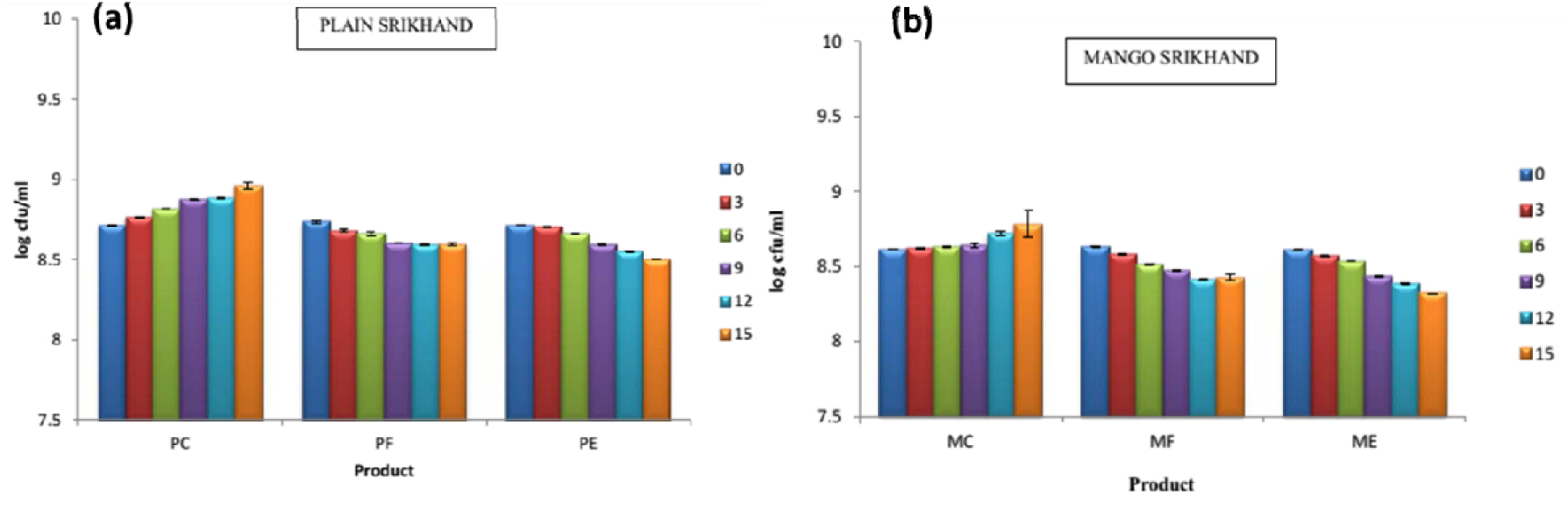
Changes in the microbial count of plain and mango srikhand during storage, *M-Mango, P-Plain, C-Control, F-Unencapsulated Free peptides, E-Encapsulated peptides

### 3.21. Sensory evaluation of srikhand

Sensory evaluation plays a crucial role in the development of any new product, as it provides insights into consumer preferences and acceptability. In this study, sensory attributes of srikhand were assessed over a two-week period to evaluate flavor, body and texture, color and appearance, and acidity. A panel of seven trained members carried out the evaluation for both plain and mango srikhand. Significant effects (P<0.05) were observed for treatment (control, free peptides, encapsulated peptides), storage days, srikhand type, and their interactions. On day 1, the overall acceptability scores for control samples were 89 ± 0.3 for plain and 91.28 ± 0.42 for mango (on a 100-point scale).

Samples with encapsulated peptides showed comparable scores (89.14 ± 0.34 for plain and 91.57 ± 0.48 for mango), whereas samples with free peptides showed a slight decline in acceptability (87 ± 0.53 for plain and 90.85 ± 0.5 for mango), likely due to the bitterness imparted by peptides. By day 5, a slight decrease in overall scores was noted for all treatments. Control samples maintained higher acceptability (87.28 ± 0.57 for plain and 87 ± 0.53 for mango), while encapsulated peptide samples showed no significant difference compared to control (84.71 ± 0.09 for plain and 86.85 ± 0.85 for mango), suggesting that encapsulation effectively masked peptide bitterness. However, a marked decline was recorded for unencapsulated peptide samples (83.42 ± 0.7 for plain and 81.85 ± 0.67 for mango) showed in figure 13. At the end of storage, control and encapsulated peptide samples retained similar acceptability scores (75.28 ± 0.4 and 74.57 ± 0.4 for plain, 77.57 ± 0.5 and 77.57 ± 0.7 for mango), whereas free peptide-treated samples showed a further decline (70.71 ± 0.7 for plain and 72.85 ± 0.45 for mango), confirming that encapsulation helped maintain sensory quality throughout storage.

**Figure 13.**
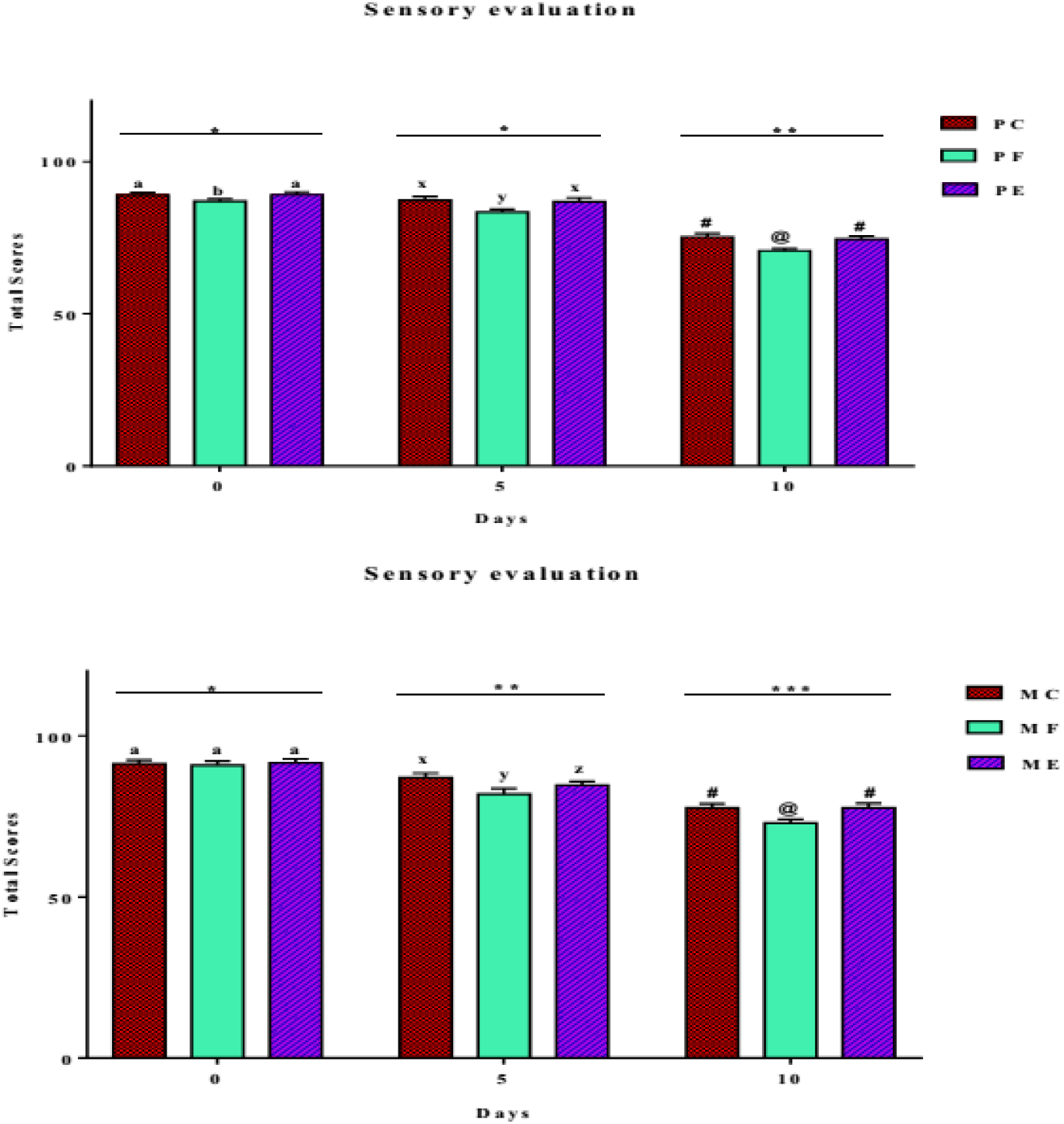
Sensory evaluation of plain and mango srikhand during storage *M-Mango, P-Plain, C-Control, F-Unencapsulated Free peptides, E-Encapsulated peptides

Overall, there was a significant decrease (*P*<0.05) in overall acceptability of the both type of srikhand with respect to days (0, 5, 10 days) may be due to increase in acidity as well as whey separation. When the type of srikhand was compared mango srikhand scored well and was more preferred as compared to plain srikhand due to addition of mango pulp which helped in masking of the bitterness of the peptides as well as the taste of double emulsion.

### 3.22. Identification of peptides from mass spectrometry (LC-MS/MS)

Peptide profiling of the <3 kDa, <5 kDa, and <10 kDa fractions of fermented sheep milk was carried out using LC-MS/MS. The resulting spectra were analyzed through Mascot and the Swiss-Prot (NCBI) database for peptide identification. A total of 3,137 peptides were detected across the three molecular weight fractions, and their amino acid sequences were confirmed through database matching.

### 3.23. *In silico* prediction of bioactive and antimicrobial peptides

A total of approximately 3,137 peptides were identified from the <10 kDa, <5 kDa, and <3 kDa fractions and confirmed through database analysis. In silico screening of antimicrobial potential was performed (Figure 14). Peptides with a PeptideRanker score between 0.7–0.90 were shortlisted (Table 1) and subsequently evaluated using iAMPpred and ADAM servers, selecting those with scores in the 0.8–2.0 range (Table 2). Thirty-four peptides that met these criteria were analyzed for physicochemical characteristics, including molecular weight, theoretical pI, aliphatic index (an indicator of thermostability), net charge, and hydropathicity (GRAVY) using the ExPASy ProtParam tool. Toxicity prediction using ToxinPred indicated that most peptides were non-toxic (Table 2). Among these, 16 peptides were identified as cationic AMPs, while the remaining 16 were hydrophobic, as evidenced by positive GRAVY scores (0.331–1.74). The theoretical pI values ranged from 3.8 to 9.75, and the aliphatic index varied from 20 to 234, reflecting a broad range of thermostability. Secondary structure analysis using UCSF Chimera and Schiffer–Edmundson helical wheel modeling revealed amphipathic α-helical conformations, with hydrophobic and hydrophilic residues positioned on opposite faces. Twelve cationic AMPs were traced to β-casein and κ-casein. Six peptides (<10 kDa) were derived from β-casein (f208–219 EPVLGPVRGPFP, f207–219 EPVLGPVRGPFPI, f124–134 MPFPKYPVEPF, f207–219 QEPVLGPVRGPFP, f207–220 QEPVLGPVRGPFPI, f206–220 YQEPVLGPVRGPFPI), five peptides (<5 kDa) also originated from β-casein, and one peptide (f75–86 QFLPYPYYAKPV) was from κ-casein. These peptides exhibited amphipathic α-helical structures with good water solubility (Figure 15).

**Figure 14.**
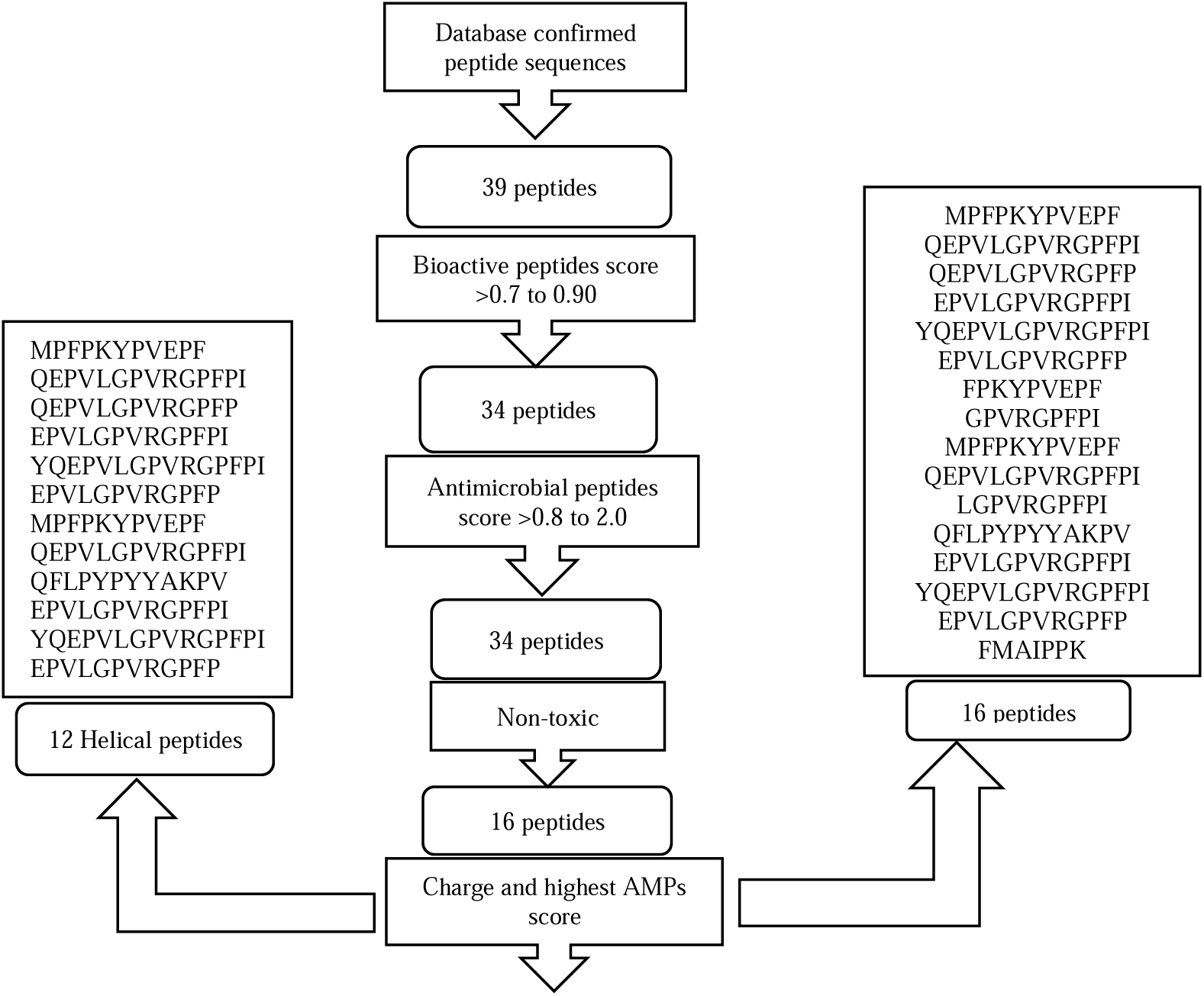
Screening process for AMPs peptides based on bioactivity, helical structure, charge, antimicrobial score, and toxicity.

**Figure 15.**
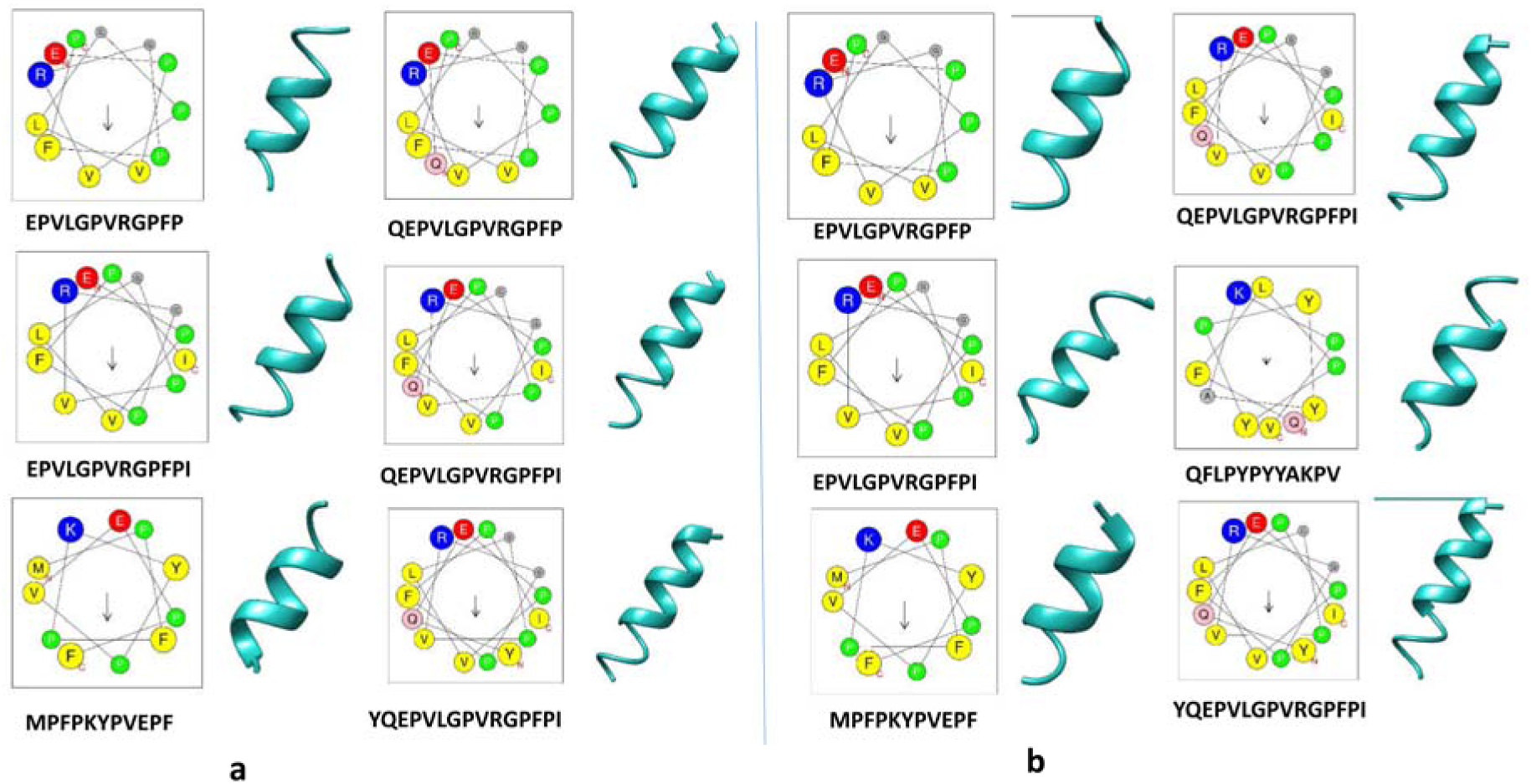
The Helical wheel projections and three-dimensional molecular modelling structure of <10kDa and <5kDa peptides of fermented sheep milk. The sheep milk derived peptide residues are number starting from the N terminus. Hydrophobic and positive charge residues were defined with yellow and blue colour, respectively. (a) <10kDa and (b) <5kDa three-dimensional molecular model of peptides was predicted by UCSF Chimera.

**Table 1.**
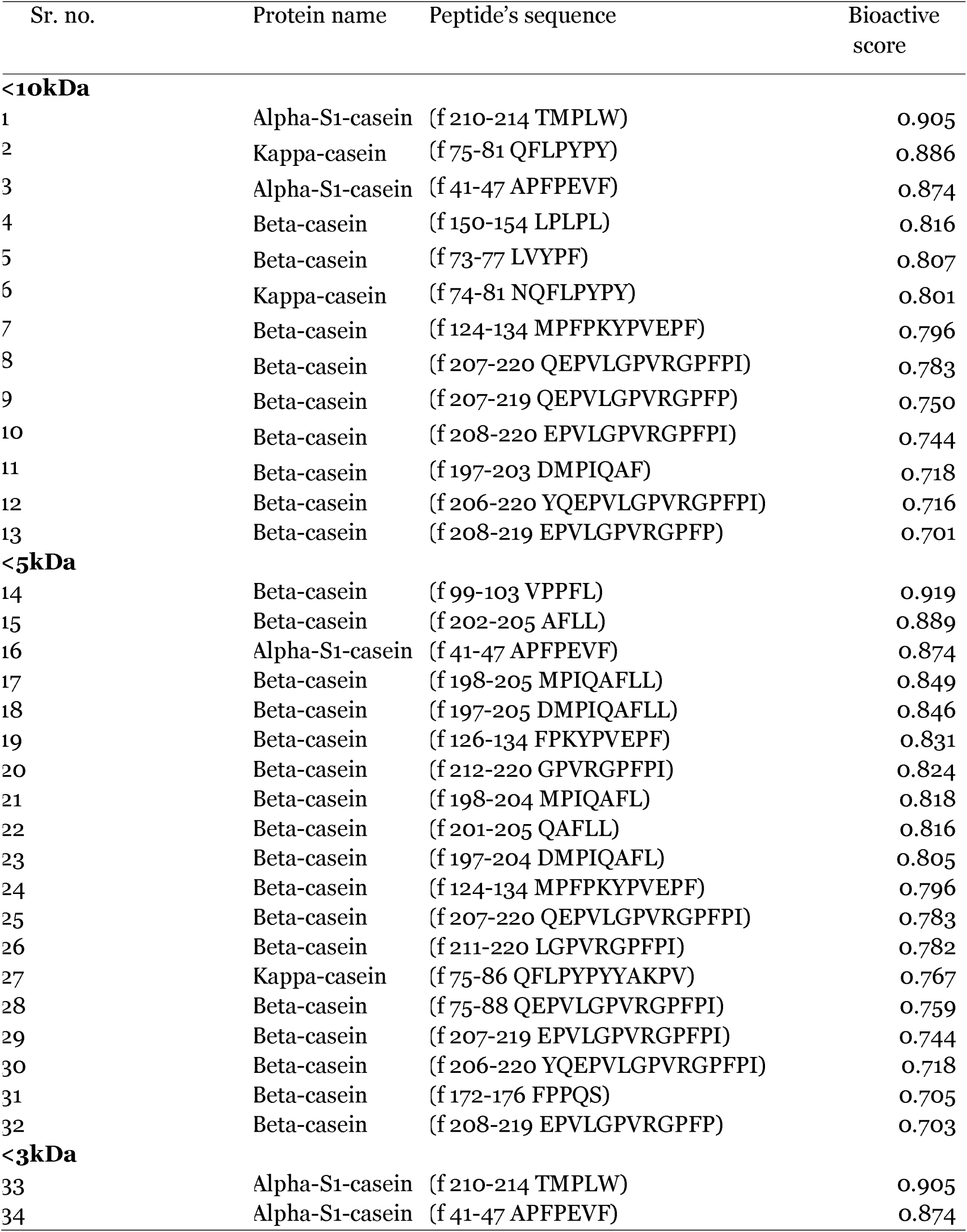
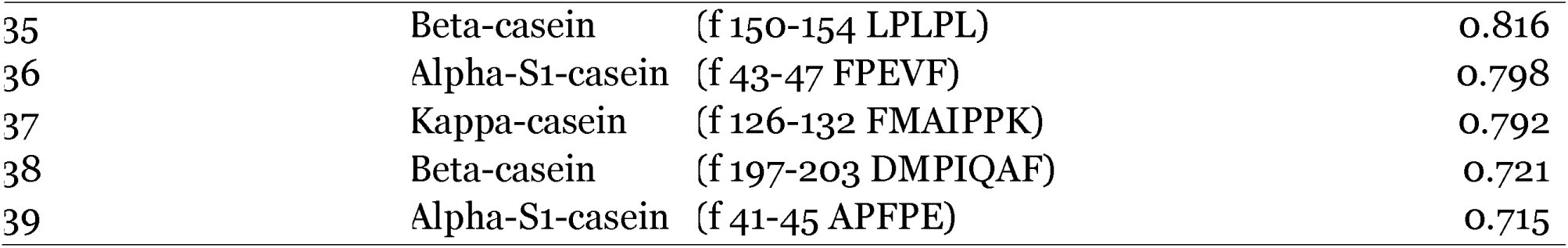
Selected bioactive peptides potential score and protein fragments name of peptide sequences derived from the fermented sheep milk proteins.

**Table 2.**
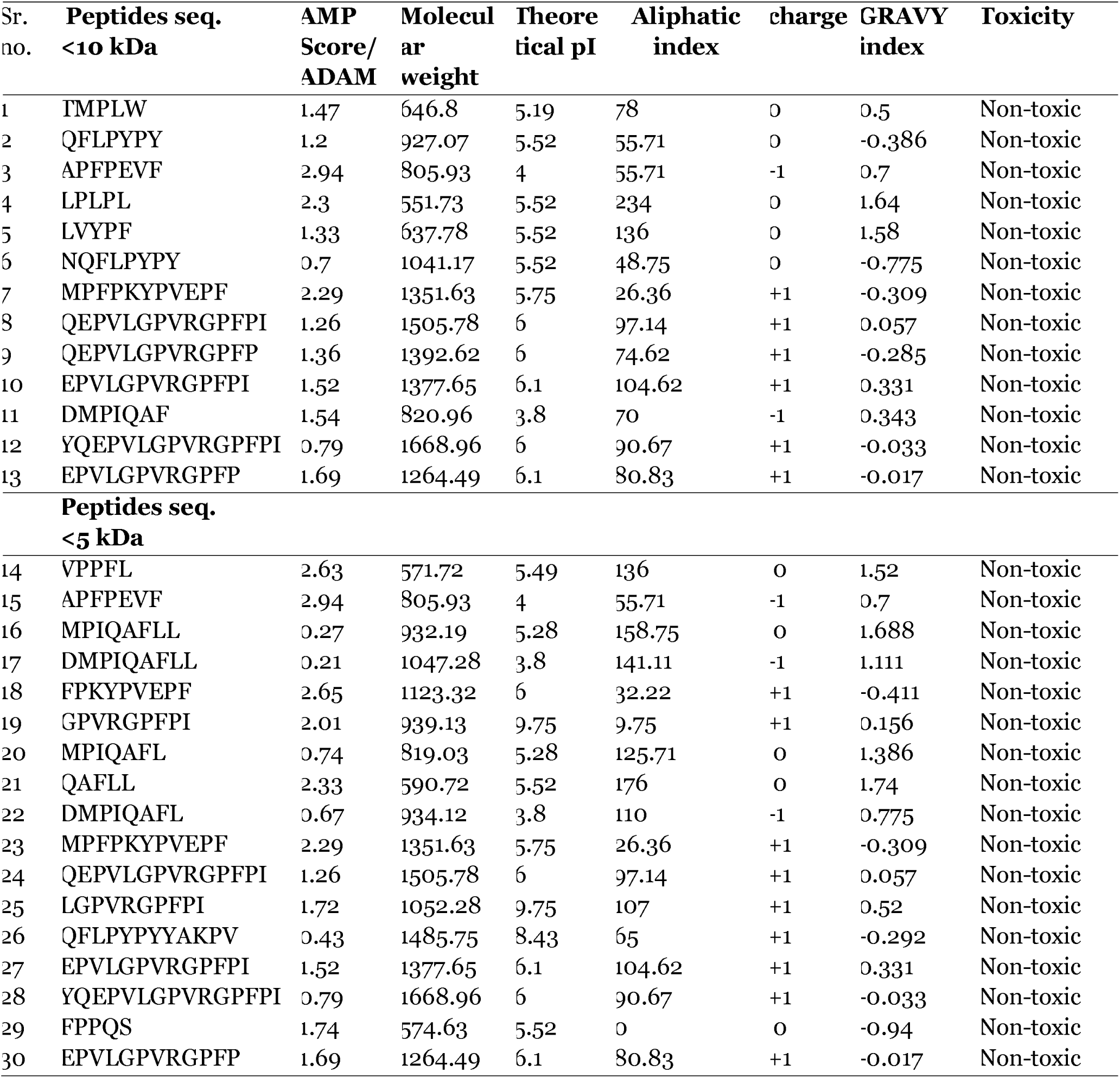
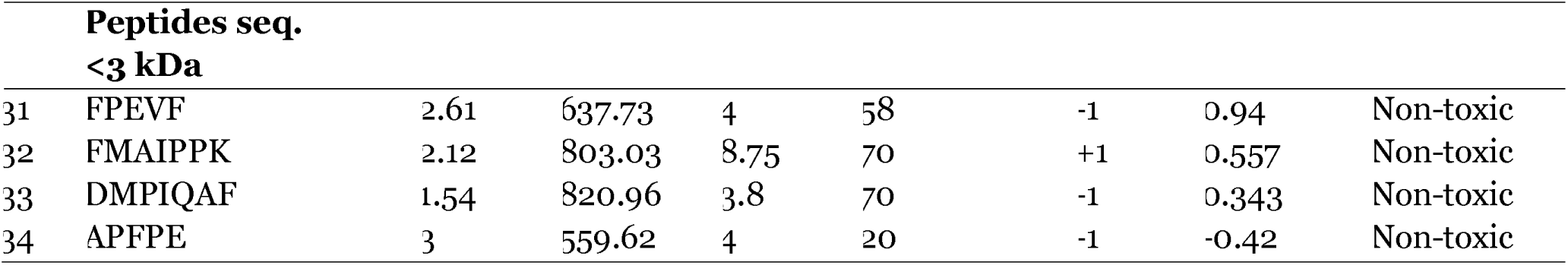
Selected AMPs peptide sequences derived from the fermented sheep milk < 10 kDa, <5kDa and <3kDa fraction, based on charge, molecular weight and highest antimicrobial potential score.

### 3.24. Peptide profiling of fermented sheep milk with *Lactobacillus rhamnosus C25*

*In silico* analysis of peptide sequences derived from LC-MS/MS profiling of Lactobacillus rhamnosus C25–fermented sheep milk revealed that the identified peptides were distributed across nearly all major milk proteins (Table 1). A total of 28 antimicrobial peptides (AMPs) were mapped to β-casein, with nine belonging to the <10 kDa fraction, seventeen to the <5 kDa fraction, and two to the <3 kDa fraction. Additionally, seven AMPs were traced to αs1-casein, including two from the <10 kDa fraction, one from the <5 kDa fraction, and four from the <3 kDa fraction. Furthermore, three AMPs were derived from κ-casein, with one peptide identified in each of the <10, <5, and <3 kDa fractions (Figure 16).

**Figure 16.**
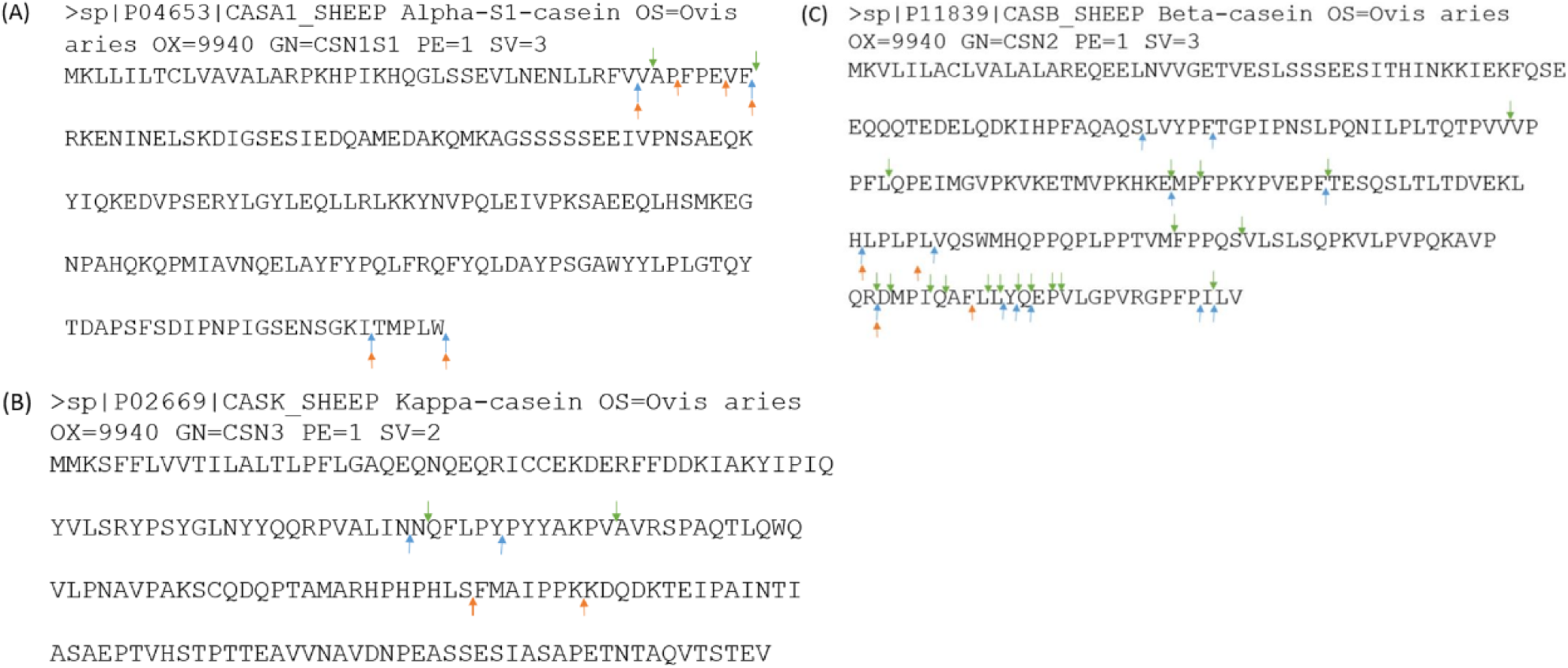
Location peptides in (A) α-s1 casein, (B) κ-casein and (C) β-casein proteins of sheep milk proteolysis by lactobacillus rhamnosus C25. Regions of highest proteolytic activity are demonstrated by arrow in different colour, the milk derived peptides fractions are represented by different arrow colour, where blue represents <10 kDa, green represents <5 kDa and red represents <3 kDa peptides.

## Conclusions

The present study demonstrates that sheep milk fermented with proteolytic Lactobacillus rhamnosus C25 is a rich source of bioactive peptides exhibiting antioxidant and antimicrobial activities. These peptides hold promise as functional ingredients for food applications. Initially, microencapsulation was performed using a two-fluid nozzle air atomization technique and incorporated into flavoured milk; however, limited dispersion of microcapsules prompted the use of an alternative encapsulation strategy. Subsequently, peptides were encapsulated via a double emulsion (W/O/W) system and added to plain and mango-flavoured srikhand. Evaluation of biofunctional activities over a two-week refrigerated storage period revealed that both types of srikhand maintained stable antioxidant and antimicrobial properties. These findings suggest that sheep milk– derived peptide fractions have strong potential as functional food ingredients, capable of enhancing the nutritional and health-promoting qualities of various food and beverage products.

## Author’s contribution

S.M., & S.V., contributed to conceptualization of study. D. I., S.M., & M.S.S., contributed to experiments, data collections and formal analysis of data. S.V., provided the facilities for the HPLC & other instrumentation. D.I., S.M., & M.S.S., wrote the manuscript. K.G., & A.P., contributed in editing and drafting the manuscript. All authors reviewed the manuscript.

## Data availability

Data will be made available on request.

## Acknowledgement

The authors express thanks towards ICAR - National Dairy Research Institute for providing the infrastructure and fund to carry out this study, and experimental dairy of the ICAR-National Dairy Research Institute, Karnal, Haryana for their cooperation during the paneer whey collection.

## References

1. Schanbacher, F. L., Talhouk, R. S., Murray, F. A., Gherman, L. I., & Willett, L. B. 1998). Milk-borne bioactive peptides. International Dairy Journal, 8(5-6):393–403.

2. Korhonen, H., & Pihlanto-Leppälä, A. (2003). Milk–derived bioactive peptides: Formation and prospects for health promotion. Handbook of Functional Dairy Products, 109–124.

3. Gobbetti, M., Minervini, F., & Rizzello, C. G. (2007). Bioactive peptides in dairy products. Handbook of Food Products Manufacturing, 489–517.

4. Haenlein, G. F., & Wendorff, W. L. (2006). Sheep milk. Handbook of Milk of Nonbovine Mammals, 137–194

5. Haenlein, G. F. W. (2001). Past, present, and future perspectives of small ruminant dairy research. Journal of Dairy Science, 84(9): 2097–2115.

6. Pulina, G., Nudda, A., Battacone, G., & Cannas, A. (2006). Effects of nutrition on the contents of fat, protein, somatic cells, aromatic compounds, and undesirable substances in sheep milk. Animal Feed Science and Technology, 131(3):255–291.

7. Moatsou, G., & Moschopoulou, E. (2014). Microbiology of raw milk. Dairy Microbiology and Biochemistry, 1–38.

8. Mayer, H. K., & Fiechter, G. (2012). Physical and chemical characteristics of sheep and goat milk in Austria. International Dairy Journal, 24(2), 57–63.

9. Geerlings, A., Villar, I. C., Zarco, F. H., Sánchez, M., Vera, R., Gomez, A. Z., & Duarte, J. (2006). Identification and characterization of novel angiotensinconverting enzyme inhibitors obtained from goat milk. Journal of Dairy Science, 89(9):3326–3335.

10. Kamau, S. M., Cheison, S. C., Chen, W., Liu, X. M., & Lu, R. R. (2010). AlphaϑLactalbumin: Its Production Technologies and Bioactive Peptides. Comprehensive Reviews in Food Science and Food Safety, 9(2):197–212.

11. Korhonen, H. (2009). Milk-derived bioactive peptides: From science to applications. Journal of Functional Foods, 1(2):177–187.

12. Zhang, L., Li, J., & Zhou, K. (2010). Chelating and radical scavenging activities of soy protein hydrolysates prepared from microbial proteases and their effect on meat lipid peroxidation. Bioresource Technology, 101(7):2084–2089.

13. Singh, B. P., Vij, S., & Hati, S. (2014). Functional significance of bioactive peptides derived from soybean. Peptides, 54:171–179.

14. Singh, M. N., Hemant, K. S. Y., Ram, M., & Shivakumar, H. G. (2010). Microencapsulation: A promising technique for controlled drug delivery. Research in Pharmaceutical Sciences, 5(2): 65.

15. Ma, G. (2014). Microencapsulation of protein drugs for drug delivery: strategy, preparation, and applications. Journal of Controlled Release, 193: 324–340.

16. Re, R., Pellegrini, N., Proteggente, A., Pannala, A., Yang, M., & Rice-Evans, C. (1999). Antioxidant activity applying an improved ABTS radical cation decolorization assay. Free Radical Biology and Medicine, 26(9):1231–1237.

17. Sharma, N., Purwar, N., & Gupta, P. C. (2015). Microspheres as drug carriers for controlled drug delivery: A review. International Journal of Pharmaceutical Sciences and Research, 6(11):4579.

18. Vincenzetti, S., Polidori, P., Mariani, P., Cammertoni, N., Fantuz, F., & Vita, A. (2008). Donkey’s milk protein fractions characterization. Food Chemistry, 106(2), 640–649.

19. Najafian, L., & Babji, A. S. (2014). Production of bioactive peptides using enzymatic hydrolysis and identification antioxidative peptides from patin (Pangasius sutchi) sarcoplasmic protein hydolysate. Journal of functional foods, 9, 280–289.

20. Meher, P. K., Sahu, T. K., & Rao, A. R. (2016). Prediction of donor splice sites using random forest with a new sequence encoding approach. BioData mining, 9(1), 1–25.

21. Yu, Zhiqiang, Quan Xu, Chenbo Dong, Su S. Lee, Liqian Gao, Yiwen Li, Mathew D’Ortenzio, and Jun Wu. “Self-assembling peptide nanofibrous hydrogel as a versatile drug delivery platform.” Current pharmaceutical d Tagliazucchi, esign 21, no. 29 (2015): 4342–4354.

22. Gasteiger, E., Hoogland, C., Gattiker, A., Duvaud, S., Wilkins, M. R., Appel, R. D., & Bairoch, A. (2016). Protein identification and analysis tools on the ExPASy server. 2005. The proteomics protocols handbook, 571–607.

23. Gupta, S., Kapoor, P., Chaudhary, K., Gautam, A., Kumar, R., Raghava, G. P., & Open-Source Drug Discovery Consortium. (2013). In silico approach for predicting toxicity of peptides and proteins. PloS one, 8(9), e73957.

24. Gautier, R., Douguet, D., Antonny, B., & Drin, G. (2008). HELIQUEST: a web server to screen sequences with specific α-helical properties. Bioinformatics, 24(18), 2101–2102.

25. Pettersen, E. F., Goddard, T. D., Huang, C. C., Couch, G. S., Greenblatt, D. M., Meng, E. C., & Ferrin, T. E. (2004). UCSF Chimera—a visualization system for exploratory research and analysis. Journal of computational chemistry, 25(13), 1605–1612.

26. Ribeiro, S. M. R., Queiroz, J. H., de Queiroz, M. E. L. R., Campos, F. M., & Sant’Ana, H. M. P. (2007). Antioxidant in mango (Mangifera indica L.) pulp. Plant Foods for Human Nutrition, 62(1):13–17.

27. Benichou, A, Aserin, A, & Garti, N. (2004). Double emulsions stabilized with hybrids of natural polymers for entrapment and slow release of active matters. Advances in Colloid and Interface Science, 108:29–41.

28. Cofrades, S., Antoniou, I., Solas, M. T., Herrero, A. M., & Jiménez-Colmenero, F. (2013). Preparation and impact of multiple (water-in-oil-in-water) emulsions in meat systems. Food Chemistry, 141(1):338–346.

29. Garti, N. (1997). Double emulsions—scope, limitations and new achievements. Colloids and Surfaces A: Physicochemical and Engineering Aspects, 123:233–246.

30. Sapei, L., Naqvi, M. A., & Rousseau, D. (2012). Stability and release properties of double emulsions for food applications. Food Hydrocolloids, 27(2):316–323.

31. Jenssen, H., Hamill, P., & Hancock, R. E. (2006). Peptide antimicrobial agents. Clinical Microbiology Reviews, 19(3):491–511.

32. Albenzio, M, Santillo, A, Caroprese, M, della Malva, A, & Marino, R (2017). Bioactive peptides in animal food products. Foods, 6(5): 35.

33. Champagne, C. P., & Fustier, P. (2007). Microencapsulation for the improved delivery of bioactive compounds into foods. Current Opinion in Biotechnology, 18(2):184–190.

34. Pellegrini, A. (2003). Antimicrobial peptides from food proteins. Current Pharmaceutical Design, 9(16):1225–1238.

35. Balasubramanian, A, Lee, D. S, Chikindas, M. L, & Yam, K. L (2011). Effect of nisin’s controlled release on microbial growth as modeled for Micrococcus luteus. Probiotics and Antimicrobial Proteins, 3(2):113–118.

36. Goyal, C. (2013). Purification and characterization of broad spectrum bacteriocin produced by a selected strain of lactococcus sp. and its microencapasulation. (Master dissertation, NDRI, Karnal).

37. Silva, S. V., Pihlanto, A., & Malcata, F. X. (2006). Bioactive peptides in ovine and caprine cheese like systems prepared with proteases from Cynara cardunculus. Journal of Dairy Science, 89(9):3336–3344.

38. Gómez-Ruiz, J. Á., López-Expósito, I., Pihlanto, A., Ramos, M., & Recio, I. (2008). Antioxidant activity of ovine casein hydrolysates: identification of active peptides by HPLC–MS/MS. European Food Research and Technology, 227(4):1061–1067.

39. Corrêa, A. P. F., Daroit, D. J., Coelho, J., Meira, S. M., Lopes, F. C., Segalin, J., & Brandelli, A. (2011). Antioxidant, antihypertensive and antimicrobial properties of ovine milk caseinate hydrolyzed with a microbial protease. Journal of the Science of Food and Agriculture, 91(12):2247–2254.

40. Lopez-Exposito, I., Minervini, F., Amigo, L., & Recio, I. (2006). Identification of antibacterial peptides from bovine κ-casein. Journal of Food Protection, 69(12): 2992–2997.

41. Power, O., Nongonierma, A. B., Jakeman, P., & FitzGerald, R. J. (2014). Food protein hydrolysates as a source of dipeptidyl peptidase IV inhibitory peptides for the management of type 2 diabetes. Proceedings of the Nutrition Society, 73(1):34–46.

42. Kim, S. K., & Wijesekara, I. (2010). Development and biological activities of marinederived bioactive peptides: A review. Journal of Functional Foods, 2(1):1–9.

43. Matsuzaki, K. (1999). Why and how are peptide–lipid interactions utilized for selfdefense? Magainins and tachyplesins as archetypes. Biochimica et Biophysica Acta (BBA)-Biomembranes, 1462(1):1–10.

44. Benkerroum, N. (2010). Antimicrobial peptides generated from milk proteins: a survey and prospects for application in the food industry.International Journal of Dairy Technology, 63(3): 320–338.

45. Benkerroum, N., Boughdadi, A., Bennani, N., & Hidane, K. (2003). Microbiological quality assessment of Moroccan camel’s milk and identification of predominating lactic acid bacteria. World Journal of Microbiology and Biotechnology, 19(6): 645–648.

46. Houser, B. A., Donaldson, S. C., Kehoe, S. I., Heinrichs, A. J., & Jayarao, B. M. (2008). A survey of bacteriological quality and the occurrence of Salmonella in raw bovine colostrum. Foodborne pathogens and disease, 5(6), 853–858.

